# A single H2A variant prevents genome instability through piRNAs biogenesis and replication stress control

**DOI:** 10.64898/2026.05.19.726330

**Authors:** Anahi Molla-Herman, Maud Ginestet, Virginie Boucherit, Emilie Brasset, Clément Carré, Jean-René Huynh

## Abstract

Transposable elements (TEs) pose a major threat to genome integrity in the germline, where the piRNA pathway ensures heritable TEs silencing. How the chromatin environment that enables piRNA biogenesis is first established during oogenesis remains unclear. Here, we identify the histone variant His2Av, the single H2A variant in *Drosophila* combining H2A.Z (transcriptional) and H2A.X (DNA repair) features, as a critical regulator of the earliest steps of piRNA pathway activation. Germline-specific depletion of His2Av disrupts transcription of piRNA pathway genes, abolishes dual-strand piRNA cluster transcription, and triggers strong TEs derepression. These defects are associated with loss of Rhino recruitment despite intact H3K9me3, suggesting that His2Av contributes to the establishment of a specialized heterochromatin permissive for noncanonical piRNA cluster transcription. His2Av depletion also causes DNA damage, replication stress, and activation of Chk2- and Claspin-dependent checkpoints, leading to oogenesis arrest. Remarkably, overexpression of RNase H, but not a catalytic-dead variant, robustly rescues oocyte development, suggesting that replication stress is a major source of DNA damage in His2Av mutants. Finally, using a separation-of-function approach with a C-terminal truncation inhibiting H2A.X-like activity, we show that the essential germline role of His2Av is transcriptional (H2A.Z-like). Together, our findings reveal that His2Av primes germline chromatin for piRNA pathway initiation while limiting transcription–replication conflicts during early oogenesis.

## INTRODUCTION

Eukaryotic chromatin is built from nucleosomes, the fundamental units of genome packaging, each consisting of two copies of the canonical histones H2A, H2B, H3, and H4 wrapped by DNA (Kornberg and Lorch 1999; Luger et al. 2012). These core histones are among the most conserved proteins in eukaryotes, reflecting their essential roles in stabilizing chromatin structure, regulating accessibility, and supporting processes such as transcription, replication, and DNA repair. Canonical histones are produced predominantly during S phase and deposited onto newly replicated DNA through dedicated histone chaperones, ensuring faithful propagation of chromatin architecture across cell division (Martire and Banaszynski 2020; Wong and Tremethick 2025; Talbert and Henikoff 2017). Their N-terminal tails are targets for a rich repertoire of post-translational modifications, such as acetylation, methylation, phosphorylation, and ubiquitination, that dynamically modulate nucleosome stability and recruit effector proteins to modify gene expression and maintain genome integrity (Millán-Zambrano et al. 2022). For example, histones acetylation is generally associated with actively transcribing regions (Verdin and Ott 2015; Hebbes et al. 1988; Talbert and Henikoff 2021) whereas the trimethylation of H3 on lysine 9 (H3K9me3) is associated with silenced heterochromatin (Lachner et al. 2001; Bannister et al. 2001; Janssen et al. 2018; Padeken et al. 2022; Grewal 2023).

In addition to these replication-coupled histones, most eukaryotes encode a diverse collection of histone variants, which are deposited independently of DNA replication and often impart specialized structural or regulatory features to chromatin. Histone variants differ from canonical histones by only a few amino acids. Yet, these small changes can confer profound effects on nucleosome dynamics, transcriptional competence, and genome stability (Martire and Banaszynski 2020; Wong and Tremethick 2025; Talbert and Henikoff 2017; Luger et al. 2012). For example, the H2A family alone includes variants implicated in transcriptional activation (H2A.Z), DNA repair signaling (H2A.X), and chromatin compaction (macroH2A). Although many histone variants have well-defined biochemical properties, their biological functions *in vivo* remain less clear, especially regarding whether they possess tissue-specific or developmental-stage-specific roles, like H2A.B in testis and H2A.W in plants (Martire and Banaszynski 2020). Germ cells, which undergo dramatic chromatin remodeling and face unique genome integrity challenges, such as meiotic recombination and protection from transposable elements (TEs), are likely to depend on specialized chromatin environments that histone variants may help establish (Molla-Herman et al. 2014; Pang et al. 2023). However, the specific contributions of individual histone variants to germline chromatin regulation remains incompletely understood.

The *Drosophila* ovary provides a powerful system in which to address these questions. Oogenesis begins in the germarium, a spatially organized structure at the anterior tip of each ovariole, where germline stem cells (GSCs) reside. GSCs divide asymmetrically to produce a new GSC and a cystoblast, which undergoes four mitotic divisions with incomplete cytokinesis to generate 16-cell cysts. These cysts progress in an ordered manner through region 1, where mitotic divisions occur. In region 2a, cysts enter meiosis and initiate the specialized transcriptional program of early oogenesis. In region 2b, one cell can be identified as the oocyte, while the remaining 15 cells adopt the nurse-cell fate. As cysts exit the germarium into region 3, they become surrounded by somatic follicle cells and progress through the subsequent stages of oocyte growth and patterning (Huynh and St Johnston 2004; Hughes et al. 2018). The germarium therefore shows the earliest chromatin transitions of germline differentiation, including mitosis, the onset of meiosis, the establishment of heterochromatin domains and the transcriptional activation of pathways required for genome defense (Molla-Herman et al. 2014; Hughes et al. 2018; Pang et al. 2023).

Transposable elements (TEs) represent a unique and important threat faced by germline cells. TEs survival depends on their transmission to the next generation, being particularly active in the germline (Senti and Brennecke 2010; Iwasaki et al. 2015; Ozata et al. 2019; Onishi et al. 2021). Their mobilization can disrupt gene expression and generate DNA breaks, which can activate DNA damage checkpoint proteins such as Chk2 (*mnk*/*loki* in *Drosophila*), leading to atrophied ovaries and sterility, thus avoiding transmission of mutations to the next generation (Klattenhoff et al. 2007; Chen et al. 2007). The resulting selective pressure exerted by TEs has driven the evolution of the piRNA pathway, a small-RNA based immune system relying on small non-coding RNAs (sncRNAs), 23 to 29 nt long, targeting TEs by sequence complementarity (Brennecke et al. 2007; Czech et al. 2018; Ozata et al. 2019). In *Drosophila*, the germline piRNA pathway are associated to nucleases from the Argonaute family: Aub, Ago3 and Piwi. In the cytoplasm, piRNAs bound to Aub or Ago3 enter into an amplificatory production loop with the targeted TE mRNAs, the ping-pong cycle, in a perinuclear organelle of nurse cells, called the nuage. When bound to Aub/Ago3, piRNAs can induce Post-Transcriptional Gene Silencing (PTGS) by targeting and cleaving TEs mRNA (Brennecke et al. 2007; Lim and Kai 2007; Senti and Brennecke 2010). piRNAs can also bind to Piwi, enter the nucleus and induce Transcriptional Gene Silencing (TGS) of TEs (Sienski et al. 2012; Le Thomas et al. 2013). TGS is achieved by the deposition of heterochromatin marks at TEs loci: the trimethylation of H3 on Lysine 9 (H3K9me3) and the recruitment of Heterochromatin Protein 1 (HP1a). Most piRNAs are derived from precursor transcripts produced by discrete genomic loci known as piRNA clusters. Germline clusters are enriched for TEs fragments and are predominantly dual-strand, producing piRNAs from both genomic strands. Dual-strand cluster transcription relies on an unusual heterochromatin state, marked by H3K9me3 and recognized by the Rhino-Deadlock-Cutoff (RDC) complex, which promotes noncanonical transcription and delivers precursor transcripts to the piRNA biogenesis machinery (Klattenhoff et al. 2009; Malone et al. 2009; Mohn et al. 2014; Ozata et al. 2019). Interestingly, although Piwi is broadly nuclear in the soma and the germline, it is transiently absent from germline nuclei in region 1, a region known as the piwi-less pocket (pilp) (Dufourt et al. 2014). This transient absence of Piwi has been interpreted in different ways: either as a window during which piRNA-mediated transcriptional silencing is minimal, or as a period marking the initiation of piRNA biogenesis, before TGS via Piwi occurs. Whether this region represents a chromatin environment permissive to piRNA cluster activation, TEs transcription, or a transition point for assembling the piRNA machinery has remained an open question.

A fundamental challenge in understanding germline genome defense is therefore to determine when and how the piRNA machinery is first established during oogenesis, what chromatin features license dual-strand cluster transcription, and how these processes are coordinated with the onset of meiosis. The identity of the chromatin factors that define piRNA-producing loci, and how they interact with DNA-damage checkpoints, remains unclear. In particular, the earliest transcriptional events in the germarium have been difficult to dissect, as they occur prior to the massive amplification of piRNAs observed later in oogenesis.

Histone variants and their contributions to TEs silencing have not been extensively studied. In the mouse embryo, the histone chaperone CAF-1 was shown to mediate the replacement of the H3 variant H3.3 with the variants H3.1/H3.2 at retrotransposon loci (Hatanaka et al. 2015). This replacement is associated with the deposition of repressive histone marks and contributes to TEs silencing. Similarly, we previously showed that in *Drosophila* ovaries mutant for CAF-1, TEs are upregulated, leading to Chk2-dependent DNA damage checkpoint activation (Clémot et al. 2018). Interestingly, it has been shown that *D. melanogaster* linker histone dH1 is required for transposon silencing and to preserve genome integrity in wing discs (Vujatovic et al. 2012). Moreover, depletion of H1 in *Drosophila* Ovarian Somatic Cells (OSCs), derepresses many TEs and their surrounding genes by increasing the chromatin accessibility of these loci, without affecting the density of H3K9me3 and HP1a (Iwasaki et al. 2016). In *Drosophila*, His2Av was also identified in genetic screens for TE repression (Czech et al. 2013; Handler et al. 2013). His2Av is the single H2A variant in flies and carries structural and functional motifs corresponding to both H2A.Z and H2A.X (Baldi and Becker 2013). However, the precise contribution of His2Av to piRNA pathway initiation has not been explored in detail, and it remains unclear whether its essential germline functions are primarily transcriptional, structural, or related to DNA damage response (DDR).

In this study, we sought to address these fundamental questions by focusing specifically on a crucial germline developmental window: the earliest steps of oogenesis, when germline cysts transition from mitotic divisions into meiosis. Our work identifies the earliest chromatin requirements for piRNA pathway activation and reveals that the single histone variant His2Av plays a critical role in initiating germline TEs silencing at the onset of oogenesis.

## RESULTS

### 1) His2Av is required for germline transposon silencing

We performed a small-scale genetic screen to identify factors required for silencing a transposon-based fluorescent reporter during early *Drosophila* oogenesis, specifically within the germarium (see Material and Methods). To this end, we expressed a collection of shRNAs in early germline cells using the *nanos*-*Gal4* driver (Rørth 1998). We screened 146 lines and we focused on one shRNA targeting the histone variant His2Av, because it had also been independently recovered in two previous screens for components required for TE repression (Czech et al. 2013; Handler et al. 2013), although its function in this context had not been analyzed further. Interestingly, knockdown of His2Av, specifically in the germline by using *nanos-Gal4* (*nos-Gal4*) driver, leads to female sterility and the formation of atrophied ovaries, arrested in early stages (st.4-5) (Fig. 1A, B). We confirmed the specificity of this shRNA for His2Av (Fig. S1A).

**Figure 1:**
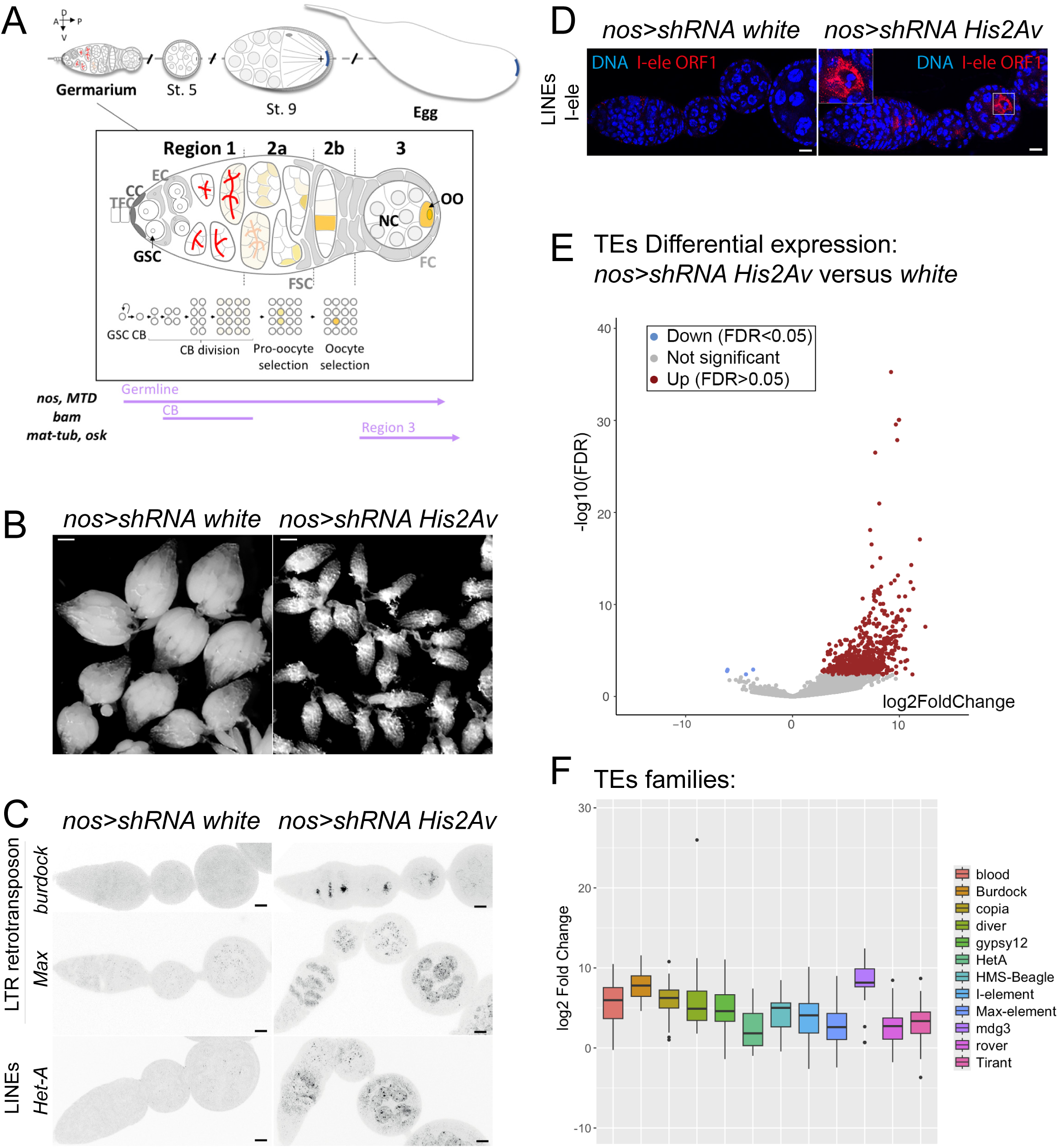
His2Av is required for TEs silencing and oogenesis development in *Drosophila*. **A.** Scheme of the main steps of *Drosophila* oogenesis, germarium organization and developmental regions. Bottom: violet arrows represent the expression profiles of different germline drivers used in this study. **B.** Ovary size of *nos>shRNA white* and *His2Av* germline mutant flies raised at 25°C. **C.** RNA FISH against transcripts from *burdock*, *Max* and *Het-A* TEs in control ovaries *nos>shRNA white* and *His2Av* germline mutants. **D.** I-element ORF1 immunostaining in control *nos>shRNA white* and *His2Av* germline mutant ovaries. A magnification image shows the oocyte, with I-ele accumulation in *nos>shRNA His2Av* mutant ovaries. Scale bar: 10 µm. **E.** Volcano plot showing TEs differential expression in *nos>shRNA His2Av* germline mutant ovaries versus control *nos>shRNA white* ovaries, after DESeq2 analysis. Colored dots correspond to |log2FC|>2, FDR<0.05. Grey dots, not significant. **F.** Distribution of log2 Fold Change values of major TEs families in *nos>shRNA His2Av* germline mutants versus virgin *nos>shRNA white* ovaries.

We validated the requirement for His2Av in transposable-element silencing by RNA FISH using probes against multiple retrotransposon families, including LTR and LINE elements. Germline knockdown of His2Av resulted in a strong transcriptional derepression of *Het-A*, *Max*, and *burdock* (Fig. 1C) and a mild expression of *copia, HMS-Beagle* and *gypsy-12* (Fig. S1B). In addition, the I-element ORF1p protein was translated and specifically accumulated in the oocyte of His2Av-depleted germaria (Fig. 1D). Importantly, we obtained identical results with a different genetic background, by doing germline clones (GLCs) with a null allele of *His2Av^810^* (Fig. S1C).

To define the temporal window during which His2Av is required for TEs silencing, we used distinct Gal4 drivers to deplete His2Av in different regions of the germarium (Molla-Herman et al. 2014; Hudson and Cooley 2014). Expression of the His2Av shRNA using *MTD-Gal4*, active throughout early oogenesis similarly to *nos-Gal4*, also resulted in small ovaries arrested around stage 5, I-ele ORF1 expression (Fig. 1A and S1D) and females were completely sterile. Knockdown using *bam-Gal4*, which is active in mitotically dividing cysts in region 1, had no detectable effect on TEs silencing or ovarian morphology (Fig. 1A and S1D). Similarly, depletion of His2Av beginning in region 3 using *mat-Gal4* or *oskar-Gal4* produced no phenotype, and these females were fertile (Fig. 1A and S1D). These findings indicate that His2Av is required likely in GSCs and in region 2a-2b of the germarium, when the 16-cell cyst enters meiosis.

To obtain a global view of transposable-element derepression, we performed RNA-seq on germline mutant ovaries (*nos*>sh-*His2Av*) and compared them with stage-matched virgin control ovaries (*nos*>sh-*white*). This analysis revealed widespread upregulation of transposable elements across multiple classes and families, as observed by RNA FISH (Fig. 1E, F).

Together, these results demonstrate that the histone variant His2Av is essential for the silencing of transposable elements when germline cysts enter meiosis in region 2a of the germarium.

### 2) His2Av loss disrupts dual-strand piRNA cluster expression and piRNA pathway proteins

The strong transcriptional derepression of transposable elements in His2Av-depleted ovaries suggested a defect in the germline piRNA pathway. TEs repression relies mainly on PIWI-clade proteins and on small piRNAs derived from piRNA clusters, and siRNA or tRNA fragments (Ozata et al. 2019; Onishi et al. 2021; Schorn and Martienssen 2018). We first examined the expression of proteins of the piRNA pathway. Analysis of our transcriptome dataset revealed that most genes encoding germline piRNA pathway proteins were down-regulated in His2Av-depleted ovaries, with the notable exception of *piwi* (Fig. 2A). Consistent with this, immunofluorescence analysis showed reduced expression of Aubergine, Ago3 and Mael, as well as a failure of these proteins to localize to the nuage, a perinuclear structure essential for piRNA biogenesis (Fig. 2B and S2A). In contrast, Piwi remained nuclear and its protein levels appeared largely unaffected (Fig. 2B).

**Figure 2:**
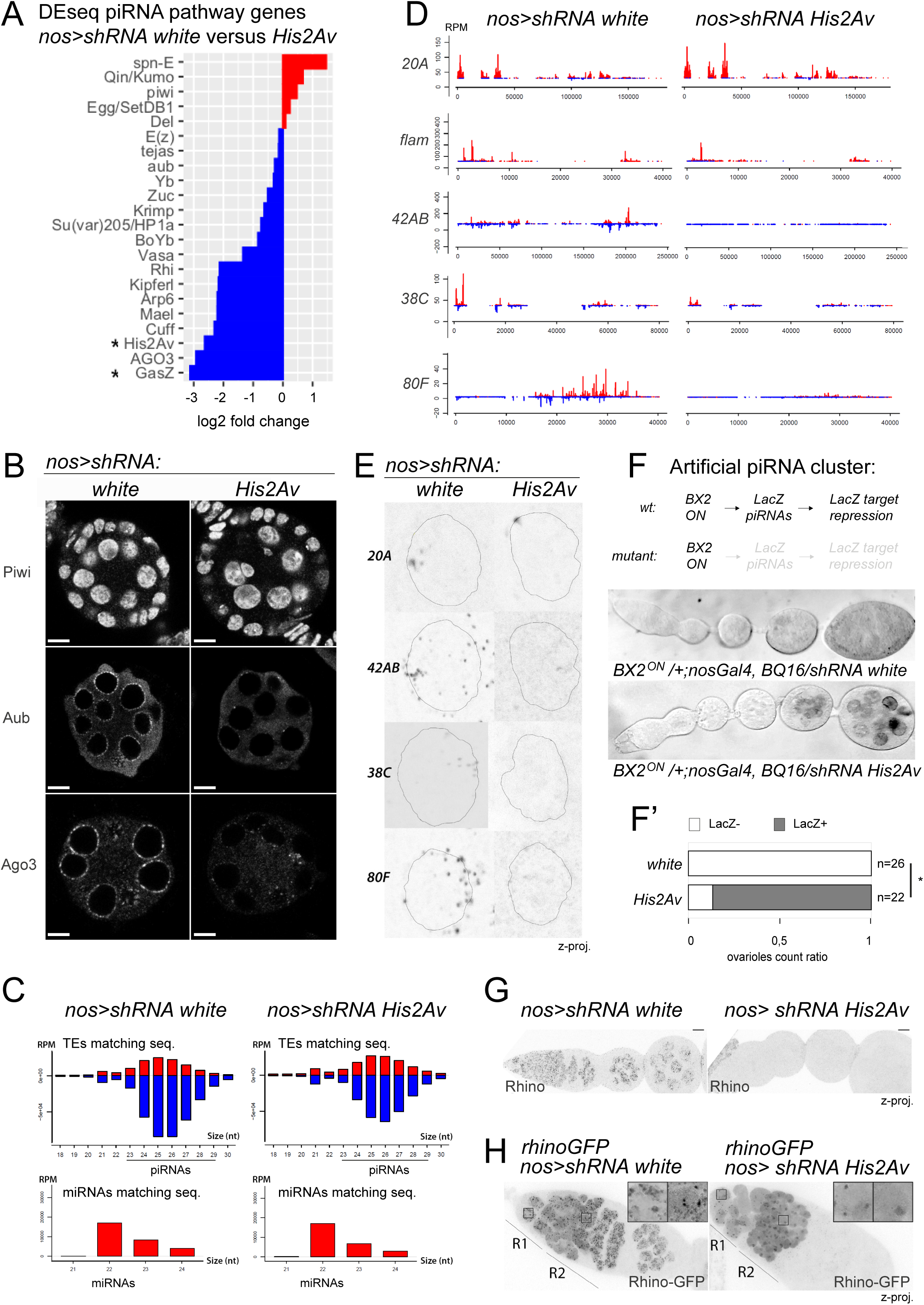
His2Av plays an important role in the piRNA pathway by regulating the transcription of some piRNA genes and the expression of dual piRNA clusters. **A.** Differential expression of major piRNA pathway genes in *nos>shRNA His2Av* germline mutant ovaries versus virgin *nos>shRNA white*. Star: significantly deregulated genes. **B.** Immunostaining of PIWI proteins (Piwi, Aub, Ago3) in *nos>shRNA white* and *His2Av* germline mutant ovaries. Scale bar: 10 µm. **C.** Size histograms of sncRNAs matching TEs (1 mismatch, multimappers randomly mapped) in virgin *nos>shRNA white and His2Av* germline mutants. Reads are normalized in RPM. piRNAs and miRNA sequences are highlighted. **D.** Multimapping piRNA reads (23-29 nt) on piRNA clusters (defined in Brennecke et al. 2007) in RPM are shown in virgin *nos>shRNA white and His2Av* germline mutants. **E.** Z maximum intensity projections of piRNA clusters transcripts, obtained by RNA FISH in *nos>shRNA white* and *His2Av germline* mutant ovaries. Nuclear profile corresponding to DNA staining is depicted with a thin black line. **F.** Artificial piRNA cluster assay: β-Gal activity assay in control *nos>shRNA white* and *His2Av* germline mutants, which received an active LacZ piRNA-producing-*BX2* cluster from their mother. piRNAs silence a LacZ target BQ16 in *wt* conditions. In mutant conditions with piRNA alteration, β-Gal staining is observed (dark signal). **F’.** Count ratio of ovarioles with β-Gal staining in *nos>shRNA white* and *His2Av* germline mutants. Data were compared by a chi^2^ test. *: p-value=6.623e-09. **G.** Z maximum intensity projection of endogenous Rhino immunostaining. Scale bar: 10 µm. **H.** *rhinoGFP* over-expression in *nos>shRNA white* and *His2Av* germline mutants. Insets show Region 1 and Region 2 Rhino-GFP signal. Z proj, Z maximum intensity projection.

To assess the small-RNA component, we sequenced ovarian small RNAs from *His2Av* germline mutants. Total piRNA abundance was reduced but not abolished (Fig. 2C), and remaining piRNAs retained a canonical ping-pong signature (Fig. S2B), indicating that processing remained functional. However, piRNAs from germline dual-strand clusters were selectively lost. Major clusters such as *42AB*, *38C*, and *80F* were nearly depleted, whereas uni-strand clusters *20A* and somatic clusters (*flamenco*) were unaffected (Fig. 2D and S2C, D).

To determine whether piRNA cluster transcription was impaired, we performed RNA FISH on individual loci. Dual-strand clusters (*42AB, 38C, 80F*) failed to be transcribed in *His2Av* knockdowns, whereas the 20A uni-strand cluster remained active (Fig.2E). Satellite-DNA loci normally expressed in the germline (1.688) were also silenced in *His2Av* mutant conditions (Fig. S2E). An artificial dual-strand cluster inserted at the BX2 locus similarly failed to generate LacZ-piRNAs in *His2Av* mutants, as indicated by derepression of its target, a β-gal BQ16 reporter (Casier et al. 2019) (Fig. 2F and F’). Finally, we observed that germline mutant males for *His2Av* were subfertile, and had a slight stellate crystal formation phenotype, as typically found in *aubergin^HN2/N11^* mutants (Fig. S2F). Indeed, piRNA pathway mutant males are known to be sterile or subfertile, and contain crystalline aggregates of Stellate-coded protein, since piRNAs/rasiRNAs *Suppressor of Stellate* from the Y chromosome are altered, and cannot silence the X-linked *Stellate* repeated genes (Aravin et al. 2004; Molla Herman and Brasset 2021).

These results show that His2Av loss disrupts the piRNA pathway at two levels: loss of key piRNA pathway proteins and a specific failure to transcribe germline dual-strand piRNA clusters, leading to depletion of germline piRNAs required for transposon silencing.

### 3) His2Av regulates germline heterochromatin and piRNA cluster expression

His2Av could contribute to piRNA cluster transcription either by directly facilitating recruitment of the transcriptional machinery or indirectly by inducing chromatin modifications required for piRNA cluster activation, such as H3K9me3, H3K27me3 and Rhino recruitment. Immunostaining revealed that Rhino protein was almost completely absent in *His2Av* germline mutants (Fig. 2G), consistent with a twofold reduction in *rhino* mRNA levels (Fig. 2A). However, overexpression of GFP-tagged Rhino failed to rescue Rhino recruitment to chromatin (Fig. 2H) and ovary growth (Fig. S3A), indicating that His2Av is also required for Rhino binding to chromatin. In addition, loss of His2Av did not impair heterochromatin assembly globally, as H3K9me3 and H3K27me3 levels were not altered (Fig. S3B and S3C). Together, these results indicate that *His2Av* is required for recruiting piRNA-activating factors such as Rhino.

Because transcriptional silencing of transposable elements correlates more strongly with HP1 binding than with H3K9me3 levels (Le Thomas et al. 2013), we examined HP1 localization using an HP1-RFP transgene. In control ovaries, HP1-RFP accumulated in the germarium and nurse cells. However, in *His2Av* germline mutants, HP1-RFP was not able to rescue the oogenesis arrest and was barely detectable in the germarium, indicating a delay in heterochromatin establishment during early oogenesis (Fig. S3A, D).

Altogether, these findings demonstrate that His2Av plays an important role in establishing a heterochromatic state permissive for piRNA cluster expression during early germline development. However, the severely reduced ovary size observed in *His2Av* germline mutants suggests additional His2Av-dependent functions in oogenesis.

### 4) His2Av controls the expression of genes important for early germline differentiation

In *His2Av* germline mutants, oogenesis is arrested around stage 5, a stage characterized by the distinctive five-lobed polytene chromosomes morphology in nurse cell nuclei (Fig. 3A). To determine whether global nuclear organization was perturbed, we visualized the three major *Drosophila* chromosomes using oligopaint DNA FISH (Nguyen and Joyce 2019). Mutant and control nurse cell nuclei displayed comparable chromosome domains, indicating that global nuclear organization remained intact in the absence of His2Av (Fig. 3B). We next assessed whether general transcriptional activity was impaired by examining RNA polymerase II phosphorylation on Ser5. Active transcription was readily detectable in *His2Av* germline mutants, with no obvious reduction relative to controls. We noticed occasional additional Pol II foci in the germarium that may correspond to ectopic TEs transcription (Fig. 3C).

**Figure 3:**
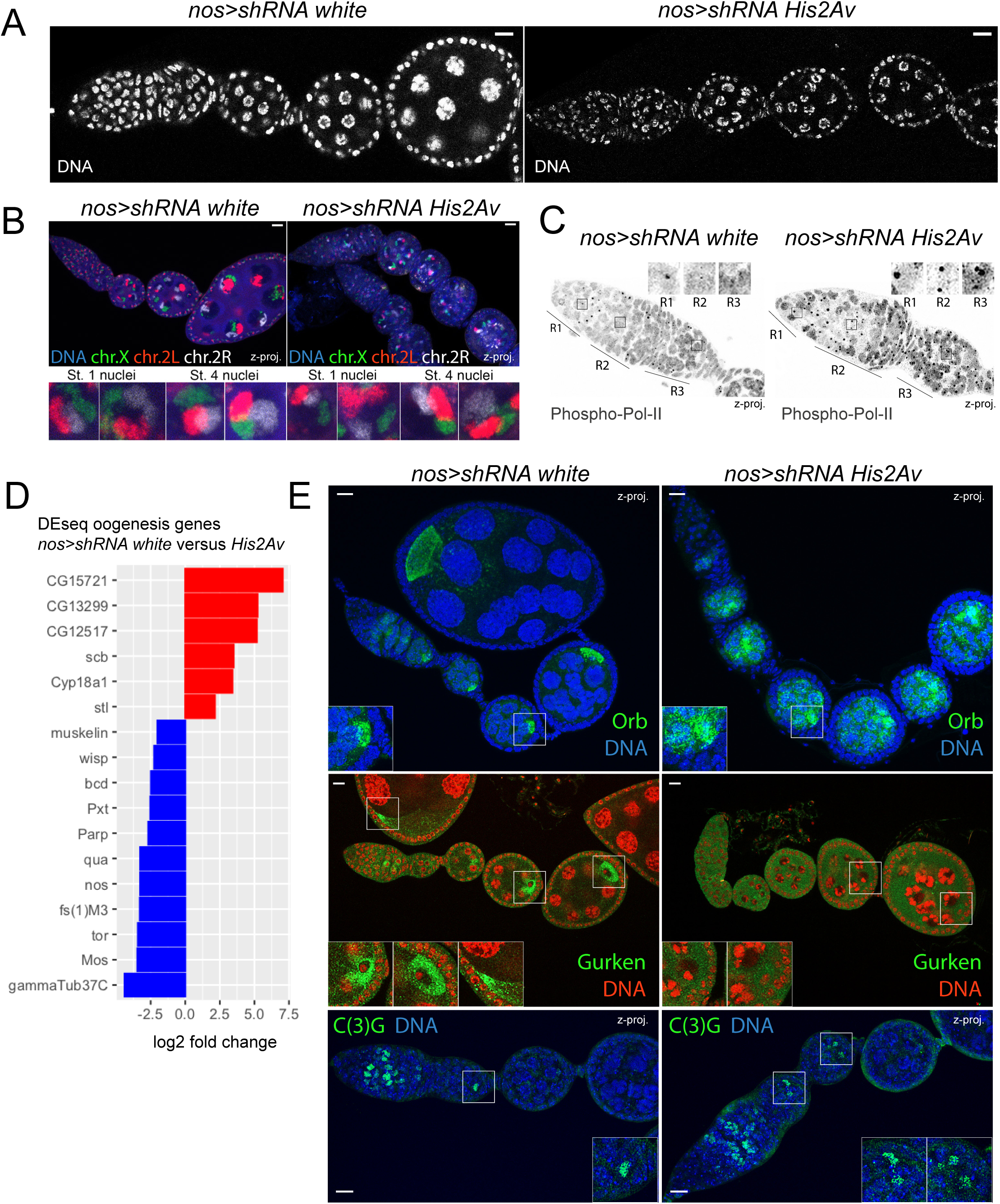
*His2Av* germline mutant flies accumulate oogenesis defects. **A.** DNA staining of *nos>shRNA white* and *His2Av* germline mutants. Scale bar: 10 μm. **B.** DNA FISH Oligopaint showing chromosome X, 2L and 2R territories in *nos>shRNA white* and *His2Av* germline mutant flies. Scale bar: 10 µm. Bottom: Z-proj. of stage 1 and stage 4 nurse cell nuclei. **C.** Immunostaining of phosphorylated RNA Polymerase II (Phospho-Pol-II) in *nos>shRNA white* and *His2Av* germline mutant ovaries. Insets show R1-R3 magnifications. Z proj, Z maximum intensity projections. **D.** Differential expression of major oogenesis genes in *nos>shRNA His2Av* germline mutant ovaries versus virgin *nos>shRNA white*. **E.** Immunostaining in *nos>shRNA white* and *His2Av* germline mutants for different oogenesis proteins. Top: orb oocyte marker, in green. Middle: Gurken protein, in green. DNA is shown in red to highlight nuclei. Bottom: synaptonemal complex protein C(3)G, in green. Insets show magnifications of the oocyte in control and mutant ovarioles. Z proj, Z maximum intensity projection. Scale bar: 10 µm.

Transcriptome analysis identified 249 significantly upregulated and 106 significantly downregulated genes (|logFC|>1.5, FDR<0.05) in mutant ovaries; many involved in developmental processes (Fig. S4A). Notably, key regulators of oogenesis such as *nanos* and *wispy* were significantly downregulated (Fig. 3D). Gene set enrichment analysis (GSEA) revealed that “non-coding RNA metabolic process” was among the most strongly downregulated categories, consistent with impaired piRNA pathway gene expression. Additional downregulated pathways included those related to fertilization, embryonic polarization, and mitotic/meiotic cell-cycle regulation, potentially explaining the developmental arrest (Fig. S4B).

To identify the early oogenesis defects, we analyzed markers of oocyte differentiation. In *wild type* germaria, Orb progressively accumulates into a single cell, the oocyte, in region 2b (Lantz et al. 1994) (Fig.3E). In contrast, in *His2Av* germline mutants, Orb remained in several cells, indicating a delay in the restriction of the oocyte fate into a single cell (Fig. 3E). In addition, localization of the oocyte dorsal-ventral determinant Gurken was abolished, suggesting DNA damage checkpoint activation (Ghabrial and Schüpbach 1999; Riechmann and Ephrussi 2001; Abdu et al. 2002) (Fig. 3E). We next monitored meiotic progression by tracking the inner synaptonemal complex component C(3)G, which starts to form in all germline cells before becoming restricted to a single cell in *wild type* germaria (Page and Hawley 2001; Mehrotra and McKim 2005). Instead, in *His2Av*-depleted cysts, two cells retained synaptonemal complex C(3)G in region 3, with one showing reduced intensity (Fig. 3E). This indicates incomplete reversion of the second pro-oocyte to the nurse-cell fate. A delay in synaptonemal complex restriction is a hallmark of early DNA damage checkpoint activation in region 2a/2b (Joyce and McKim 2010).

Overall, the absence of Gurken protein and delayed synaptonemal complex restriction suggest the activation of an early checkpoint in *His2Av* germline mutants, consistent with the early defects in piRNA pathway function described above.

### 5) His2Av prevents replication stress and DNA-damage checkpoint activation

To determine whether His2Av depletion induces DNA damage, we monitored single-stranded DNA using an RPA-GFP reporter (Blythe and Wieschaus 2015), as phosphorylated-His2Av cannot be used in this context. In *His2Av* germline mutants, RPA-GFP foci were strongly increased throughout cyst differentiation, indicating significant DNA damage (Fig. 4A, B). The main effector of the DNA-damage checkpoint in *Drosophila* oogenesis is the kinase Chk2 (Klattenhoff et al. 2007; Chen et al. 2007). To assess whether Chk2 activation contributed to the developmental arrest, we combined *His2Av* germline knockdown (KD) with a heterozygous mutation in *mnk* (Chk2). Loss of a single *mnk^P6^*allele partially rescued ovary growth by ∼40%, demonstrating that Chk2 activation contributes to the oogenesis arrest of *His2Av* germline mutants (Fig. 4C, C’).

**Figure 4:**
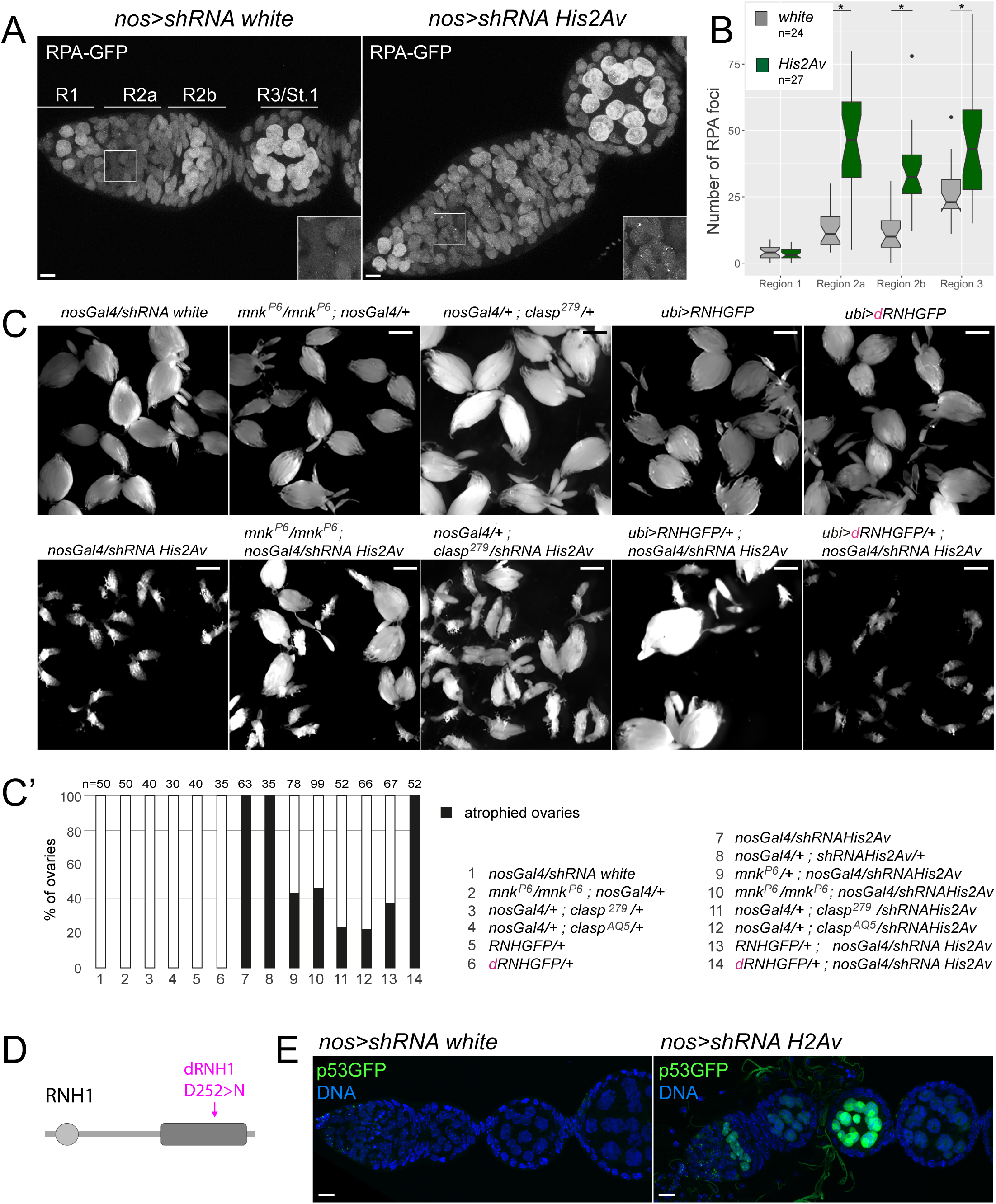
The accumulation of DNA damage from different sources leads to oogenesis arrest in *His2Av* germline mutant flies. **A.** Examples of Z maximum intensity projections of RPA-GFP in *nos>shRNA white* or *His2Av* mutant germaria and stage 1 egg chambers, used for quantification of RPA foci per region. **B.** Quantification of RPA-GFP foci in control (grey) and *nos>shRNA His2Av* (green) mutant ovaries. DNA damage levels across oogenesis development are shown (Mann-Whitney U test): Region 1 p-value=0.7966 ; Region 2A: p-value=8.632e-08 ; Region 2B: p-value=1.69e-07 ; Region 3: p-value=0.0008308. **C.** Ovary size in rescue experiments where *His2Av* germline mutation is combined with mutants of the DNA damage checkpoint pathway (*chk2/mnk* or *claspin*) or with the over-expression of RNase H (RNH-GFP) or catalytically dead RNase H (dRNH-GFP). Top panels : control conditions. Bottom panels: Rescue conditions. Scale bar: 2.5 mm. **C’.** The percentage of atrophied ovaries is shown for different genotypes. **D.** RNaseH (RNH1) domains are depicted. Catalytically dead dRNH: D252>N, in violet. **E.** p53 activity was analyzed with a p53GFP biosensor (Wylie et al. 2014) expressed in *nos>shRNA white* and *His2Av* germline mutant flies. Scale bar: 10 µm.

p53 acts downstream of Chk2 (Hirao et al. 2000). We thus examined a p53 fluorescent reporter (Wylie et al. 2014) and observed strong activation in *His2Av* germline mutants (Fig. 4E). However, we did not detect early Caspase-3 activation (Yu et al. 2002), indicating that apoptosis was not significantly induced at the time of developmental arrest. Cleaved Caspase-3 was detected only later, in mid-oogenesis (after several stages 5-arrested egg chambers), and never in early germarium regions (Fig. S5A). To determine whether cell death was responsible for the arrest, we coexpressed the apoptosis inhibitor p35 with *His2Av* shRNA in the germline. Overexpression of p35 (Mazzalupo and Cooley 2006) did not rescue the early oogenesis block, indicating that oogenesis arrest occurs independently of apoptosis (Fig. S5B, B’).

Multiple sources could underlie the DNA damage observed in His2Av-depleted germ cells, including meiotic double-strand breaks, replication stress, or activation of transposable elements. We previously showed that replication stress blocks oogenesis and activates Claspin, a key component of the replication stress specific DNA damage checkpoint (Lee et al. 2012; Molla-Herman et al. 2015). We thus generated double mutants combining *His2Av* germline KD with a mutation in *claspin*. We observed a partial, but consistent, rescue of ovary growth, indicating that replication stress is one contributor to the oogenesis arrest observed in *His2Av* germline mutants (Fig. 4C).

One possible source of replication stress could be an excessive transcription of transposable elements (El Hage et al. 2014; Gambelli et al. 2023; Stanek et al. 2025). High levels of transposon transcription are known to induce RNA-DNA hybrids (R-loops), which in turn impede replication fork progression (García-Muse and Aguilera 2016). To examine whether R-loops contribute to the *His2Av* phenotype, we overexpressed the *Drosophila* RNase H homolog (RNH-GFP), which specifically resolves RNA-DNA hybrids (El Hage et al. 2010; García-Muse and Aguilera 2016). As a control, we generated a catalytically inactive RNase H variant (dRNH-GFP) (Chen et al. 2017) by mutating a key residue in the active site (D252>N) (Fig. 4D). Overexpression of *wild-type* RNase H strongly rescued ovarian growth in *His2Av* germline mutant females (Fig. 4C, C’). Importantly, the catalytic-dead RNase H did not rescue the oogenesis arrest (Fig. 4C, C’).

These findings indicate that replication stress accumulate in His2Av-deficient germ cells and contribute to checkpoint activation. Together with the increase of RPA foci and our genetic analyses of Chk2 and Claspin, these results suggest that the developmental arrest in *His2Av* mutant germ cells is caused by the activation of multiple DNA-damage checkpoints.

### 6) His2Av C-terminal tail is dispensable for TE silencing

His2Av is the sole H2A variant in *Drosophila melanogaster* and combines two functions carried by distinct mammalian variants: an H2A.Z-like role in transcriptional regulation and an H2A.X-like role in DNA damage repair, through phosphorylation of its C-terminal tail (Baldi and Becker 2013). To separate these activities and study which of these functions is important during early oogenesis, we used CRISPR/Cas9 to generate a ΔCter-His2Av allele lacking the final 14 amino acids of the C-terminus, including the conserved phosphorylation site required for DNA damage signaling (Clarkson et al. 1999) (Fig. 5A). Loss of the C-terminal tail was confirmed by immunostaining using an antibody that recognizes the phosphorylated epitope absent in this ΔCter-H2Av fly line (Fig. S5C). Homozygous *ΔCter-His2Av* flies were viable and fertile, and their ovaries were mostly normal in size (Fig. 5B). piRNA pathway components were unaffected, as Piwi and Aubergine localization was indistinguishable from wild type (Fig. 5C). Orb localized to a single cell in region 2b, indicating that oocyte specification and polarization were intact (Fig. 5D). In contrast to *His2Av*-depleted germ cells, TEs *Max* and *gypsy12* remained fully silenced in *ΔCter-His2Av* ovaries (Fig. 5E). Consistently, I-element ORF1 was neither transcribed nor translated (Fig. 5E). These results demonstrate that the His2Av C-terminal tail is largely dispensable for piRNA-mediated TEs silencing and germline cyst differentiation.

**Figure 5:**
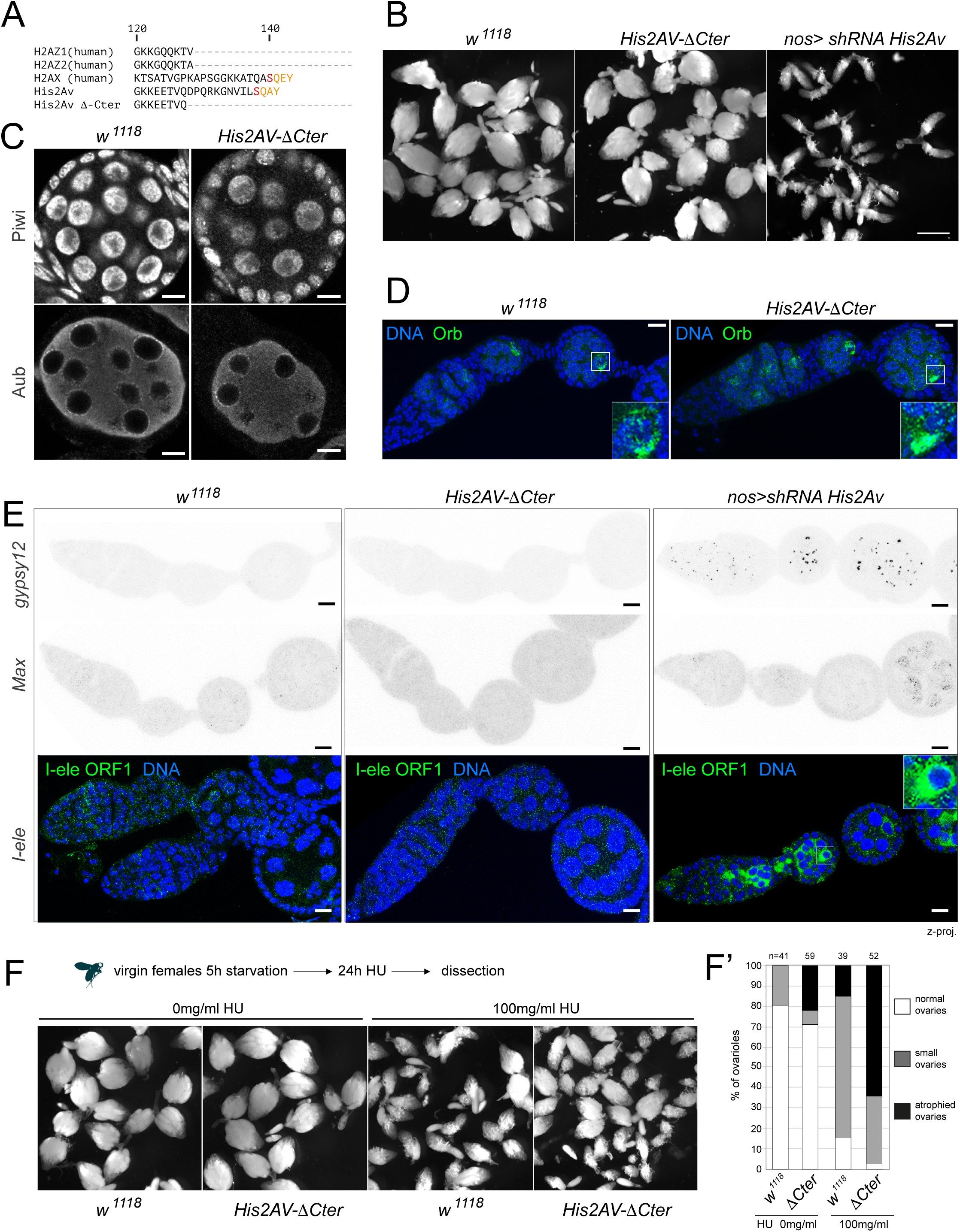
H2A.Z-like transcriptional role is required during *Drosophila* oogenesis. **A.** Protein sequence alignment (Jalview) of the two *human* H2A.Z variants (Z1 and Z2) and H2A.X, *D. melanogaster His2Av* and *His2Av-ΔC-terminal* (CRISPR-deletion of the 14 last amino acids of His2Av, containing the phosphorylation motif similar to human H2A.X). **B.** Ovary size of control *w^1118^*, *His2Av-ΔCter* homozygous flies and *nos>shRNA His2Av* germline mutants. Scale bar: 2.5 mm. **C.** Immunostaining of PIWI proteins Piwi and Aub in control flies and *His2Av-ΔCter* homozygous mutants. Scale bar: 10 µm. **D.** Immunostaining of Orb in control flies and *His2Av-ΔCter* homozygous mutants. Insets show magnifications of the oocyte. Scale bar: 10 µm. **E.** Top: RNA FISH against *gypsy12* and *Max-element* expression in control *w^1118^*, *His2Av-ΔCter* homozygous flies and shRNA *His2Av* germline mutants. Bottom: Immunostaining of *I-element* ORF1 in control, *His2Av-ΔCter* homozygous mutants and *His2Av* germline mutant ovaries. Insets show a magnification of the oocyte. Scale bar: 10 µm. **F.** Virgin control *w^1118^* and *His2Av-ΔCter* homozygous mutants flies were starved for 5h, then exposed to 0 or 100 mg/ml of Hydroxyurea (HU) for 24h, and dissected. Ovaries sizes are shown in both cases. **F’.** Quantification of atrophied ovaries (with less than 5 stage-9-egg chambers or more advances stages), small size ovaries (more than 5 stage-9-egg chambers or more advances stages) and normal ovaries.

Nevertheless, His2Av C-terminus is known to be phosphorylated in response to ionizing radiation and is required for efficient repair of irradiation-induced DSBs in *Drosophila* and mammals (Madigan 2002; Rogakou et al. 1998). Because replication stress contributes to the oogenesis arrest in *His2Av*-depleted germ cells, we tested whether the C-terminal tail was required for responding to hydroxyurea (HU)-induced replication stress (García-Muse and Aguilera 2016). Both *wild-type* and *ΔCter-His2Av* virgin females were starved for 5 hours before adding 0 mg/ml or 100 mg/ml of HU to the food. We noticed that ovary morphology was affected by starving in both *wild-type* and *ΔCter-His2Av* females, although mutant females were more affected than control flies (Fig. 5F and F’). Importantly, in the presence of HU (100 mg/ml) both *wild type* and *ΔCter-His2Av* females became sterile. However, *ΔCter-His2Av* mutants displayed a higher proportion of atrophied or small ovaries, indicating increased sensitivity to replication stress relative to controls (Fig. 5F and F’).

In *Drosophila*, the Domino-B isoform and Arp6 are core components of the major nucleosome-remodeling complex that incorporates His2Av at promoters to support transcriptional activity (H2A.Z-like) (Börner and Becker 2016; Scacchetti et al. 2020) (Fig. S6A). Consistent with previous reports, germline knockdown of Arp6 led to a strong reduction of His2Av levels in germline cells (Fig. S6C). Importantly, under these conditions, we detected expression of I-element ORF1p (Fig S6C), as well as transcripts from the retrotransposons *gypsy12* and *burdock* (Fig S6D). Moreover, combining germline *Arp6* knockdown with loss of a single *mnk^P6^* allele resulted in a strong reduction of I-element expression (Fig. S6E, F). We therefore concluded that, similarly to *His2Av* depletion, *Arp6* knockdown leads to transposon derepression and activation of the Chk2-dependent checkpoint.

Together, these findings indicate that the early oogenesis defects observed in *His2Av*-depleted germ cells arise primarily from the transcriptional function of His2Av (H2A.Z-like), rather than its function in DNA-damage response (H2A.X-like). *ΔCter-His2Av* mutant flies are however more sensitive to stress such as irradiation, starvation or HU treatment.

### 7) Ovo expression partially rescues oogenesis and TEs silencing in His2Av mutants

Our results so far indicate that the transcriptional activity of His2Av is required for transposon silencing and proper germline cyst differentiation during early oogenesis. To gain further mechanistic insights into *His2Av* mutant phenotypes, we looked for transcription factors known to regulate germline development. Interestingly, we noted a reduction in *ovo* expression, although its fold change was just below our log2 FC threshold (logFC=-0.746, FDR=0.377). Ovo was recently shown to act as a major transcriptional regulator of the germline piRNA pathway in flies, driving the expression of ∼70% of piRNA pathway genes and ∼18% of genes involved in ovarian development (Benner et al. 2024; Alizada et al. 2025). To test whether reduced *ovo* expression contributes to the defects observed in *His2Av* mutants, we introduced a GFP-tagged transgene containing the endogenous *ovo* locus (*ovo*GFP) into *His2Av*-depleted ovaries. This transgene thus provides a modest increase in Ovo levels by adding a single extra genomic copy. Importantly, this additional copy of *ovo* led to a substantial reduction in the number of atrophied ovaries in the *His2Av* mutant background, although flies remained sterile (Fig. 6A, B). Moreover, I-element protein expression was strongly decreased in these rescued ovaries, indicating that piRNA-mediated TE silencing was at least partially rescued (Fig. 6C, D). In addition, *ovoGFP* was able to partially rescue the presence of Gurken (Fig. 6E, G) and the localization of Orb into the oocyte (Fig. 6F, H). We also found that *ovoGFP* expression leads to an increase of Aub levels, and its proper localization to the cytoplasmic nuage in nurse cells (Fig. 6I).

**Figure 6.**
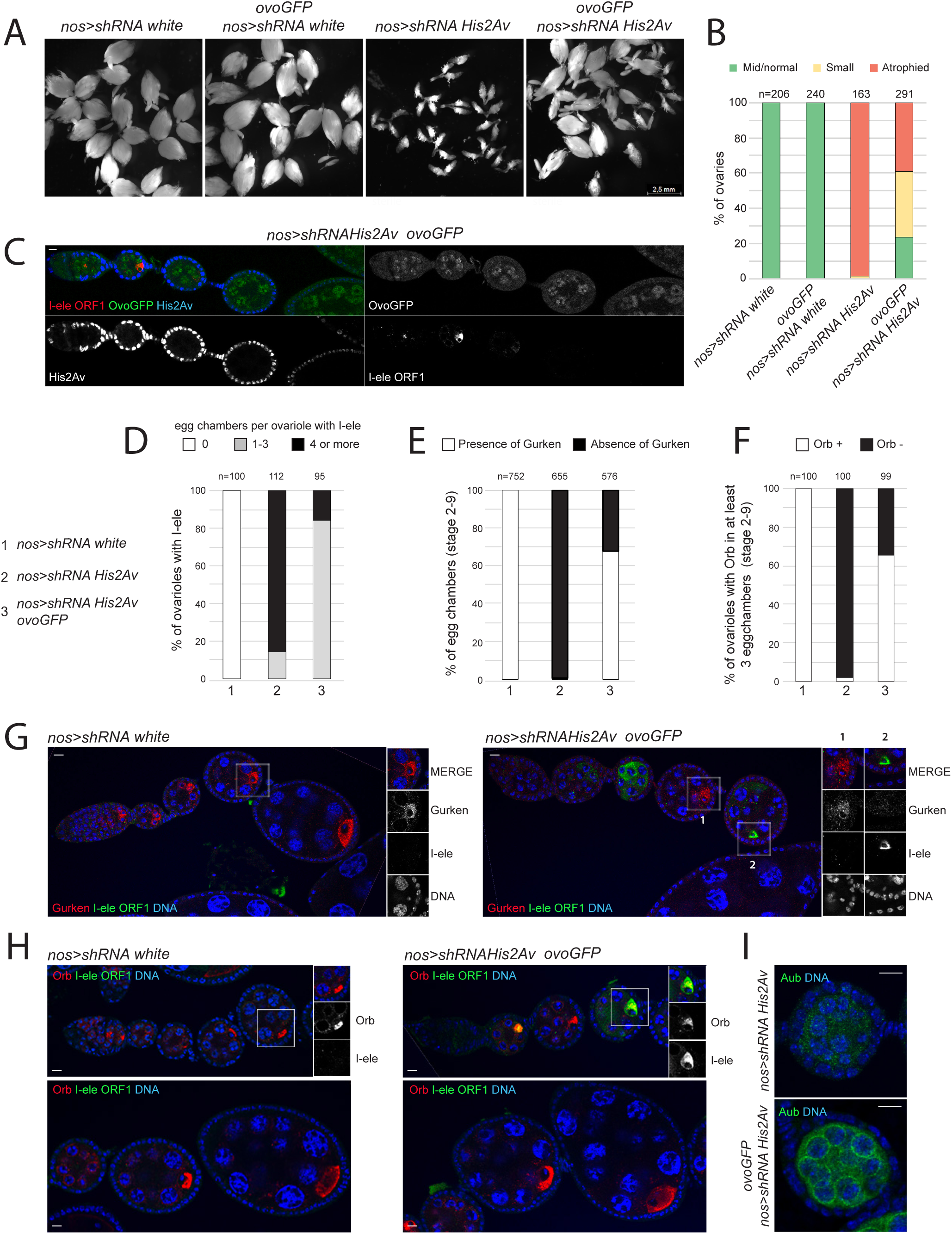
*ovoGFP* partially rescues *His2Av* mutant oogenesis arrest. **A.** Ovaries are shown in control *nos>shRNA white* and *His2Av* germline mutants, in the presence or absence of *ovoGFP* overexpression. **B.** Quantifications of the partial rescue found in A, showing: mid/normal ovaries (green), small ovaries (yellow, containing more than 5 stages-9-egg chambers or more advanced stage) and atrophied ovaries (red, with less than 5 stages-9-egg chambers or more advanced stage). **C.** Immunostaining of I-element ORF1 (red) and His2Av (blue) in *nos>shRNA His2Av* germline mutant ovaries overexpressing *ovoGFP* (green). Scale bar: 10 µm. **D.** I-ele ORF1 signal was quantified in *nos>shRNA white* and *His2Av* mutant ovaries overexpressing *ovoGFP*. Three main categories of ovarioles were found: 0 egg chambers with I-ele per ovariole. 1-3 egg chambers with I-ele per ovariole. 4 or more egg chambers with I-ele per ovariole. **E.** Quantification of the percentage of egg chambers (stage 2-9) with or without Gurken. **F.** Quantification of the percentage of ovarioles with Orb in at least 3 egg chambers (stages 2-9) with a normal Orb staining localized into the oocyte (Orb+) or with an altered diffused Orb staining (Orb-). **G.** Immunostaining for Gurken and I-ele ORF1 in control *nos>shRNA white* and *His2Av* germline mutants overexpressing *ovoGFP*. Insets show oocyte’s magnifications: 1. Example of an egg chamber rescued by *ovoGFP,* in which Gurken is expressed, without I-ele. 2. Exemple of an egg chamber not rescued by *ovoGFP*, in which Gurken is not expressed and I-ele is present. Scale bar: 10 µm. **H.** Immunostaining for Orb and I-ele ORF1 in control *nos>shRNA white* and *His2Av* germline mutants overexpressing *ovoGFP*. Insets show oocyte’s magnifications. Scale bar: 10 µm. **I.** Immunostaining for Aub in *nos>shRNA His2Av* germline mutants overexpressing *ovoGFP* or not. Insets show oocyte’s magnifications. Scale bar: 10 µm.

Together, these findings demonstrate that reduction in Ovo levels in His2Av-KD contribute significantly to both TEs transcriptional derepression and defects in early germline development.

## DISCUSSION

His2Av has been identified in at least three independent genetic screens for factors required to silence transposable elements in the *Drosophila* germline. Yet, its mechanistic contribution to this process has remained unclear. Here, we provide a comprehensive analysis demonstrating that His2Av is required for transcription of dual-strand piRNA clusters, for maintenance of piRNA pathway gene expression, and for preventing replication stress and DNA damage checkpoint activation during cyst differentiation. We further demonstrate that the C-terminal tail of His2Av, which mediates DNA-damage signaling role, is largely dispensable for piRNA-mediated silencing, whereas the transcriptional function of His2Av is essential. Finally, we show that reduced Ovo levels account substantially for the oogenesis and TEs silencing defects observed upon His2Av depletion, and that restoring Ovo expression partially rescues these phenotypes. Together, our findings establish His2Av as a central chromatin regulator that coordinates transcriptional programs, heterochromatin states, and genome stability during the earliest stages of germline development.

### 1) His2Av acts primarily through a transcriptional function reminiscent of H2A.Z in germline cells

Histone variants are required for crucial processes during animal development and are implicated in many human pathologies. Loss of H2A.Z, H3.3 and CENP-A in mouse induce gastrulation defects and embryonic lethality. They are also implicated in different human cancers and neuropathologies (Buschbeck and Hake 2017; Ghiraldini et al. 2021; Colino-Sanguino et al. 2022; Dijkwel and Tremethick 2022; Wong and Tremethick 2025). *Drosophila* His2Av is unusual among metazoan H2A variants as it combines the activities of both H2A.Z and H2A.X within a single protein. Whereas H2A.Z is broadly associated with transcriptional regulation, chromatin accessibility, and promoter activation, H2A.X is specialized for marking DNA damage via phosphorylation of its SQ motif. In *His2Av*-depleted germ cells, we detected the earliest defects in region 2a/2b, when germline cysts enter meiosis, characterized by TEs expression, RPA foci accumulation and a delay in restricting the synaptonemal complex (SC) to a single pro-oocyte. These early defects triggered the activation of checkpoint proteins and an arrest of oogenesis, later at stage 5. In contrast, ΔCter-His2Av mutant ovaries were mainly fertile, with no detectable TEs expression, and normal germline differentiation, including nuage formation. These findings align His2Av essential germline role with the transcription-associated functions of H2A.Z, rather than with the damage-signaling functions of H2A.X. Interestingly, two recent studies in mouse oogenesis indicate that the transcriptional functions of H2A.Z are also required for proper oocyte development and for maintaining genome stability. These findings raise the possibility that the transcription-associated role of H2A variants, in supporting germline development and transposon control, may be conserved across metazoans (Mei et al. 2025; Xu et al. 2025).

Our results raise the question of how His2Av, despite being a single dual-function variant, performs H2A.Z-like transcriptional functions in the *Drosophila* ovary. Our results suggest that His2Av could be required to establish a specialized chromatin environment that supports transcription of both piRNA pathway genes and germline dual-strand piRNA clusters. In particular, germline piRNA clusters require an atypical transcriptional mode that combines features traditionally associated with heterochromatic stability and euchromatic transcriptional competence. Dual-strand clusters rely on the Rhino-Deadlock-Cutoff (RDC) complex, which binds H3K9me3 and supports noncanonical transcription. However, it is not well understood why *in vivo* only a fraction of H3K9me3-decorated heterochromatin is occupied by Rhino. In this sense, it has been recently showed that Kipferl recruits Rhino to a subset of piRNA clusters in a sequence-dependent manner (Baumgartner et al. 2022). Additionally, at Kipferl-independent sites, Rhino binds to loci marked by both H3K9me3 and H3K27me3 via its chromodomain, in a specific manner (Akkouche et al. 2025). We showed that His2Av depletion eliminates Rhino without diminishing H3K9me3 or H3K27me3 levels. On the one hand, these results highlight that H3K9me3 alone is not sufficient to recruit RDC; and on the other hand, that His2Av could be a missing determinant of this noncanonical chromatin state. His2Av may therefore act upstream of RDC and Kipferl to create a chromatin landscape that permits transcription of piRNA cluster precursors.

It has been demonstrated that deleting the three main germline piRNA clusters has only modest effects on TE silencing (Gebert et al. 2021). This observation suggests that, although His2Av depletion eliminates dual-strand cluster transcription, the striking TE derepression we observe is more likely to come from a collapse in piRNA pathway gene expression rather than the loss of cluster-derived piRNAs themselves. Our transcriptome analysis identified extensive downregulation of core piRNA factors upon His2Av depletion, and one such factor, *ovo*, proved particularly important for mediating the consequences of His2Av loss. Indeed, the transcription factor Ovo was recently shown to be a master regulator of germline gene expression, required both for oocyte specification and for transcription of nearly 70% of germline piRNA pathway genes (Alizada et al. 2025) and many important oogenesis genes in the female germline (Benner et al. 2024). Although *ovo* transcripts were modestly reduced in *His2Av*-depleted ovaries, restoring *ovo* expression with a single additional genomic copy was sufficient to partially rescue ovarian morphology and significantly restore TE silencing. This could explain the loss of many piRNA pathway genes in *His2Av*-depleted cells and provide a mechanistic link between His2Av chromatin role and pathway-wide effects on TE repression. His2Av may therefore promote transcription of piRNA pathway genes at two levels: 1) by establishing a chromatin landscape that is permissive for their transcription; and 2) by promoting *ovo* transcription, which in turn activates the expression of piRNA pathway genes.

These findings raise important new questions about how His2Av regulates *ovo* expression and, more broadly, how His2Av and Ovo might function together to coordinate germline transcriptional programs. Does His2Av directly occupy regulatory elements at the *ovo* locus to promote transcription? Is His2Av directly incorporated at nucleosomes in piRNAs genes? Do His2Av and Ovo regulate together overlapping sets of piRNA pathway genes, or do they act sequentially within a hierarchical regulatory cascade? Interestingly, His2Av has been shown to act with the transcription factor Zelda to initiate transcription during zygotic genome activation in *Drosophila* embryos (Ibarra-Morales et al. 2021). By analogy, we propose that His2Av may play a similar role together with the transcription factor Ovo during germline differentiation.

### 2) His2Av loss uncovers a direct contribution of replication stress to germline DNA damage

Our results show that depletion of His2Av leads to DNA damage and activation of the Chk2-, p53-, and replication stress-responsive checkpoints, including Claspin. Faithful DNA replication is essential for the maintenance of genomic integrity; however, replication is continually challenged by DNA damage and other perturbations that induce replication stress. Such stress can arise from DNA lesions, secondary DNA structures, R-loops (DNA–RNA hybrids), nucleotide pool imbalances, highly transcribed loci (e.g., rDNA), or transposable elements. These perturbations can expose single-stranded DNA (ssDNA), generating fragile sites that predispose to double-strand breaks (DSBs) and genome instability. Cells have therefore evolved complex mechanisms to tolerate and resolve replication-associated problems. For example, R-loops can be removed by RNase H activity, and replication protein A (RPA) coats ssDNA, serving as a platform for the recruitment of checkpoint proteins. The accumulation of RPA-GFP foci throughout cyst differentiation in *His2Av* mutants, together with the partial rescue of ovarian growth upon reduction of Chk2 activity and the absence of early apoptosis, collectively demonstrate that checkpoint activation, rather than cell death, is responsible for the stage-5 oogenesis arrest. Such phenotypes are common among mutants in piRNA pathway genes (Chen et al. 2007). However, the precise causes of Chk2 activation have remained unclear. A recent study elegantly demonstrated that even a small number of DSBs by TE excision events is sufficient to trigger Chk2 activation and germ cell loss, rather than DNA damage caused by new TE insertions (Jansen et al. 2024).

Here, we provide multiple lines of evidence implicating replication stress as a major source of DNA damage in *His2Av*-depleted germ cells: 1) Claspin heterozygosity partially rescues ovarian growth; 2) TEs transcription is strongly upregulated; and 3) overexpression of RNase H, but not its catalytic-dead version, induces the most striking and complete rescue of the oogenesis arrest that we observed in this study. The most common cause of replication stress is the accumulation of R-loops. We propose that the strong TE transcription observed in His2Av-depleted ovaries could generate widespread R-loops, whose resolution by RNase H could explain the strong rescue of oogenesis. Our results therefore indicate that TE transcription, in addition to TE excision, could be sufficient to activate the DNA damage checkpoint in the germline. In this sense, it has been recently shown that R-loops are present at TEs in several species, including humans and fruit fly (Zeng et al. 2021; Paul et al. 2025; Stanek et al. 2025; Li et al. 2025). In *rhino* mutant ovaries, R-loops strongly accumulate at LTR transposon and DNA satellites (Stanek et al. 2025).

### 3) Integrating His2Av dual functionality in the germline

Our dissection of His2Av function reveals a striking separation between transcriptional and DNA-damage roles. The *ΔCter-His2Av* allele shows that, despite being the single H2A variant in *Drosophila*, modularity persists within His2Av: the N-terminal and histone-fold domains support transcriptional roles, while the C-terminal tail mediates damage signaling. This raises an interesting evolutionary question: why are these roles fused into a single variant in *Drosophila* while other species maintain their separation? One possibility is that in *Drosophila* germline cells, there is extensive transcription of repetitive elements coinciding with rapid nuclear divisions in the mitotic region of the germarium. This region is also known as the *piwiless pocket* (pilp) because of low levels of Piwi proteins and de-silencing of some transposable elements and could participate in generating genomic diversity during evolution (Dufourt et al. 2014). It may thus be advantageous for flies to have a chromatin environment with a single H2A variant, able to coordinate transcriptional regulation of these genomic regions with response to replication stress and DNA repair in these same regions.

Conversely, in species in which H2A.Z and H2A.X are encoded by distinct genes, germline cells may rely on more specialized chromatin states or checkpoint pathways to manage transcription-replication conflicts. In *Arabidopsis*, H2A.Z has been proposed to direct Ty1/copia retrotransposon insertions away from essential genes and preferentially into environmentally responsive loci. This bias may favor the generation of large-effect mutations and facilitate rapid adaptation (Quadrana et al. 2019). In yeast, H2A.Z mutants are viable but display significant growth defects under replication stress conditions, such as exposure to hydroxyurea (HU). Notably, H2A.Z is preferentially incorporated into genes required for the response to HU-induced replication stress. Consistent with these observations, we find that *ΔCter-His2Av* mutants are more sensitive to replication stress induced by HU treatment.

These results suggest that, although the C-terminal tail of His2Av is dispensable for oogenesis under normal conditions, it provides additional capacity to cope with exogenous DNA damage or replication stress. This finding further indicates that the damage-signaling, H2A.X–like function of His2Av, may become critical under environmental stress, such as HU exposure, as observed in yeast, or under conditions of pathological TEs reactivation and biased insertions, as described in *Arabidopsis*. It will therefore be of interest to determine whether His2Av genomic localization is dynamically remodeled under genotoxic stress to promote transcription of stress-induced genes, and how TEs behavior is biased in this context.

Together, our work shows that His2Av links transcriptional regulation, transposable element control, and replication stress responses in the germline. More broadly, these findings suggest that histone variants can help coordinate genome defense with developmental transcription programs, under both normal and stress conditions.

## MATERIALS and METHODS

### Fly stocks

Flies were maintained on standard medium, in incubators with a 12h light/dark cycle at 25°C, except when stated otherwise (18°C, 22°C or 29°C).

The following stocks are detailed in Table n°1.

*nosGal4,RPA-GFP* ; *nosGal4,tRFLys-GFP* and *nosGal4,GFP* flies were obtained in the laboratory by standard recombination methods.

RNH1 cDNA obtained from The FlyBi ORFeome Collection (DGRC Stock 1654593) was amplified by PCR, then cloned into the pENTR/D-TOPO Gateway vector (Invitrogen) and subcloned into the pUbi vector with a C-terminal GFP tag (Murphy and Huynh lab, DGRC). An inactive form (D252 mutated to N) was generated by Site Directed Mutagenesis (QuickChange II XL kit from Agilent). Transgenic fly lines were obtained by random insertions (BestGene).

His2Av-ΔCter flies lacking the last 14 amino acids of His2Av were generated by CRISPR/Cas9 (Well Genetics).

### Genetic Screen (Fig. S7)

The tRFs-Lys-GFP sensor was made similarly to miRNA-GFP sensor (Brennecke et al. 2005): an ubiquitous tubulin promoter, a GFP containing a NLS, and a 18nt TE sequence at the 3’UTR, complementary to a 3’CCA-tRF-Lys-TTT, known to be abundantly expressed in *wt* ovaries and altered in *Rpp30^18.2^* homozygous mutant ovaries (Molla-Herman et al. 2020). Knocking down a gene involved in tRF-mediated TEs silencing would increase or decrease GFP expression. The tRF-Lys-GFP reporter was combined with UAS/Gal4 system to perform a targeted RNAi screen, in order to identify new factors involved in tRFs gene silencing *in vivo*. *nosGal4* was recombined with the tRF-Lys reporter (both on chr. III). *nosGal4,tRFs-Lys-GFP* was crossed with *UAS::shRNA* lines of interest. UASp based shRNAs lines (valium 20, 22, 21) were chosen since they had a link with tRF biogenesis/tRNA processing/tRNA modification/tRFs binding in *Drosophila* or in other systems in the published literature. Additional lines were chosen if they were reported to be involved in TEs silencing, in the piRNA pathway or in chromatin regulation. 146 shRNA lines were screened. Two independent crosses were done per genotype. 10 ovaries per genotype were dissected per genotype. About 100 ovarioles were observed per genotype at the microscope. We found three groups based on qualitative assessment of the extent of GFP silencing in germaria: i) no effect ii) GFP overexpression in all germ cells iii) increased GFP silencing in all germ cells. *His2Av* mutants showed a complete GFP overexpression in the whole germarium.

### Flp/FRT clone generation

*FRT80-82B,His2Av^810^/TM6,Tb* flies were obtained in the laboratory with standard recombination methods. These flies were crossed with *y,w, flp; FRT82AGFP*. 3rd instar larvae were heat-shocked two times at 37°C for 2 h (morning and then afternoon), and one time the next morning for 1h. Ovaries from adult flies were dissected about 10 days after hatching. Homozygous mutant clones were recognized by the absence of GFP.

### RNA extraction and sequencing

RNA was extracted as in (Molla-Herman et al. 2015). *white* mutant virgin flies were used as a developmental control for *His2Av* mutant flies, since their ovaries are of comparable size. 20 µL of each sample (100 ng.µL^-1^) were sent to Fasteris NGS Service for sequencing. Small RNA-seq Gel Free protocol was performed on Illumina NextSeq High and paired-end mRNA-seq (polyA) on Illumina NovaSeq 6000.

### Bioinformatic Data analysis

All bioinformatic data analyses were done using R or Python softwares wrapped up Galaxy, available by the ARTbio platform (https://mississippi.sorbonne-universite.fr/login/start?redirect=%2). For with R 4.2 using R Studio.

### RNA-seq

Quality control was performed with FastQC. Reads were aligned to dm6 reference genome using STAR (default parameters, 1 mismatch). Genes and TEs were annotated separately using featureCounts and manually curated annotations from Flybase and Repbase, respectively. Reference genes annotations were downloaded on Flybase and reference TEs annotations were kindly provided by Dr. Silke Jensen (iGRED, Clermont-Ferrand), downloaded on Repbase and manually curated. All counts were pooled and analyzed with DESeq2 using default parameters. Subsequent GSEA was performed on the WebGestalt portal (https://www.webgestalt.org/), taking into account all pathways with FDR<0.01.

### Small RNA-seq

Only 18 to 30 nt reads were analyzed. Quality control was performed with FastQC. Reads were normalized to 1 million reads to allow library comparisons.

Reads mapping to reference tRNA, rRNA, miscRNA and miRNA (3 mismatches) were filtered. Remaining reads were mapped to dm6 reference genome. TE-matching 21 nt-long reads correspond to siRNAs and 23-29 nt-long to piRNAs. piRNAs in silico characterization was performed using small_rna_signatures (Antoniewski, Methods Mol.Biol. 2014) and WebLogos were generated in the eponymous web-based application (https://weblogo.threeplusone.com).

Reference piRNA clusters used are those defined in (Brennecke et al. 2007) and reference TEs sequences were downloaded on Repbase (<u>(Brennecke et al. 2007)</u>) in June 2024. Small RNA reads matching was made allowing 1 mismatch.

### Datasets Availability

Raw RNA-seq data are available in the NCBI Sequence Read Archive (SRA) under BioProject accession number PRJNA1466887.

### Hydroxyurea (HU) treatment

Flies were collected 1 day after hatching and starved for 5h in an empty tube. Then, flies were put for 24h on a medium with 0 or 100 mg.mL^-1^ HU before dissection (HU: H8627-1G Sigma).

### Immunofluorescence in ovaries and testis

Ovaries were dissected in PBS and fixed in 4% PFA for 20 minutes, except for nuage components (5 min). Then, ovaries were washed 5 times in PBS, permeabilized and blocked for 1h in PBS Triton 0.2%, BSA 3% and primary antibodies were incubated over 1 to 3 nights at 4°C. Secondary antibodies (Alexa Fluor 488, 555 or 647 conjugated antibodies from Invitrogen) were used at 1:500 dilution and incubated for 2h at room temperature, after 5 washes in PBS. Following 5 more washes, ovaries were incubated with Hoescht at 5 µg.mL^-1^ for 5 min at room temperature. Samples were then kept at 4°C overnight in mounting medium (CitiFluor^TM^ AF1, Electron Microscopy Sciences). Ovarioles were mounted on a microscope slide (Superfrost^TM^ Plus Adhesion, Epredia), removing eggs and late stages, and covered by a 18 x 18 mm, 1.5 µm-thick coverslip (VWR).

Testes were dissected in PBS and then fixed in 4% PFA in PBT 0.3% plus 3 volumes of heptane and incubated for 20 min at room temperature. After two washes in PBT 0.2% and 1 h in PBT 0.3% and BSA 3%, testes were incubated with primary antibody overnight in PBT 0.2%. After several washes, secondary antibody was added for 2 h in PBT 0.2%, testes were washed, incubated with Hoechst for 15 min in PBS, and incubated in Citifluor at 4°C until mounting.

Antibodies used in this study are detailed in Table n°2.

### RNA Fluorescence In Situ Hybridization (FISH)

Ovaries were dissected in PBS and fixed in 3.7% formaldehyde diluted in PBS for 20 min. Ovaries were washed 2 times for 10 min in RNase-free PBT (PBS, 0.3% Triton X-100) and permeabilized overnight in EtOH 70%. Then, ovaries were rehydrated in Stellaris Wash Buffer A or smRNA FISH wash buffer (2X SSC with 10% formamide) and FISH probes were incubated overnight at 37°C in Stellaris Hybridization Buffer or 2X SSC with 10% formamide, 10% dextran sulfate. 1.5 µL for TEs or 2 µL for piRNA clusters of probes were diluted in 50 µL of hybridization buffer. Samples were washed 2 times for 30 min at 37°C in Stellaris Wash Buffer A or smRNA FISH wash buffer, then incubated with Hoescht at 1 µg.mL^-1^ in the same buffer for 10 min at room temperature. Ovaries were stored at 4°C in mounting medium after a 5 min wash in Stellaris Wash Buffer B.

Except for gypsy12 and Max-element, RNA FISH probes were provided by Pr. E. Brasset (iGRED, Clermont-Ferrand). RNA FISH probes are listed in the Table n°3.

### DNA FISH Oligopainting

Oligopaints were done following Nguyen et al. 2019 (Nguyen and Joyce 2019). Ovaries were dissected in PBS and fixed 10 min in a solution made of 100 μL 4% PFA, 300μL heptane. Then, ovaries were washed 3 times quickly and 3 times 5 min in PBS 0.02% Tween20. The same washing protocol was repeated in 2X SSC, 0.01% Tween20 before incubating the ovaries for 10 min in 50% formamide. After a 3-5 h of incubation in 50% formamide at 37°C, probes and samples are denatured 30 min at 80°C. Ovaries are then incubated overnight with DNA FISH probes at 37°C. 25pmol of probes, 2R-Cy5, 2L-Cy3 and X-A488, gift from Dr. E. Joyce (University of Pennsylvania, USA), are incubated in a 40 μL reaction in 2X SSC, 0.01% Tween, 10% dextran sulfate, 50% formamide with 1 μL of RNase A. Samples are washed 2 times in 50% formamide for 30 min at 37°C, then in 20% formamide for 10 min at RT. After 4 quick washes in 2X SSC 0.01% Tween20, Hoescht at 5 μg.mL-1 is incubated for 5 min at RT. Samples are washed 2 more times in 2X SSC 0.01% Tween20 for 5 min and in 2X SSC for 10 min before removing all liquid and adding mounting medium (CitiFluorTM AF1, Electron Microscopy Sciences). Samples are kept at 4°C until mounting on a microscope slide (SuperfrostTM Plus Adhesion, Epredia). Eggs and late stages are removed and ovarioles are covered by a 18 x 18 mm, 1.5 μm-thick coverslip (VWR).

### LacZ staining

Ovaries were fixed for 10 min with glutaraldehyde 0.5%, then washed 3 times 10 min with PBS 1X. Staining was performed overnight at 37°C in PBS 1X, 1.5% X-gal (5bromo-4-chloro-3-indolyl-β-D-galactopyranoside), 3.5% ferri-ferro-cyanide. Ovaries were rinsed and stored at 4°C in mounting medium until mounting on a microscope slide as described above.

### Image acquisition

Samples were imaged using a confocal microscope (Zeiss LSM 980) and images were acquired using the ZEN software, then processed with Fiji and presented with Adobe Photoshop (v 24.5) and Illustrator (v 27.61). Z-stacks were acquired with a 0.5 µm step and are presented as Z maximum intensity projections. Same settings are always used in mutant and control conditions.

### Data analysis of images

RPA-GFP dots were counted manually in control and mutant conditions using Fiji.

For piRNA cluster transcripts RNA FISH, background was subtracted using a rolling ball of radius comprised between 20 and 50 pixels. The same setting was used for each set of probes.

### Rescue experiments

Flies were raised at 22°C and kept 1 week on yeast after hatching before dissection in PBS. Ovaries were imaged with a fluorescence stereomicroscope Leica M205 FCA using LasX software. Atrophied ovaries have less than 5 stage-9 egg chambers (or more advances stages). Small ovaries have more than 5 stage-9 egg chambers (or more advanced stages) and were counted as rescued. Normal ovaries have full developed egg chambers.

## ACKNOWLEDGMENTS

We are grateful to Antoine Boivin (Sorbonne Uni., Paris) for LacZ experiments, to Abdou Akkouche (UCA) for RNA FISH experiments, to Eric Joyce and Nguyen Son (Uni. Pennsylvania, USA) for oligopaint protocol and reagents, to Bloomington Stock Center, DSHB Hybridoma Center, for antibodies and flies. We acknowledge Silke Jensen (UCA), Julie Autaa (Sorbonne Uni., Paris) and Aurélie Tessandier (Institut Curie, Paris) for their help with bioinformatic analysis. We are grateful to the Orion Imaging facility at CIRB (Collège de France). We thank Geneviève Almouzni, Raphael Margueron and Daniel Holoch (Institut Curie, Paris) for scientific discussions; and Katalin Fejes Toth (CalTech, USA) for sharing data prior to publication. We also thank the Huynh lab for helpful comments on the manuscript.

JRH, EB and CC labs share common ANR grants (ANR-13-BSV2-0007-02 PlasTiSiPi; ANR-21-CE12-0022, BioPic). In addition, JRH lab is supported by CNRS, Inserm, Collège de France, and the Bettencourt-Schueller foundation (FBS). M.G. had fellowships from Ministère de l’Enseignement Supérieur et de la Recherche and Fondation pour la Recherche Médicale (FRM). E.B. lab is also supported by ANR JCJC CHApiTRE ANR-20-CE12-0005. C.C. lab is also supported by ANR-24-CE17-7018-01 and Fondation pour la Recherche sur le Cancer ARCPJA2022070005319.

## AUTHOR CONTRIBUTIONS

A.M.-H. and J.-R.H. conceived the study and wrote the manuscript. A.M.-H. and M.G. performed and analyzed experiments. V.B. performed cloning experiments. E.B. and C.C. provided methods and reagents. A.M.-H., C.C. and J.-R.H. supervised the study. J.-R.H. provided funding, laboratory space and resources. All authors commented on the manuscript.

---

**Supplementary Figure 1.**
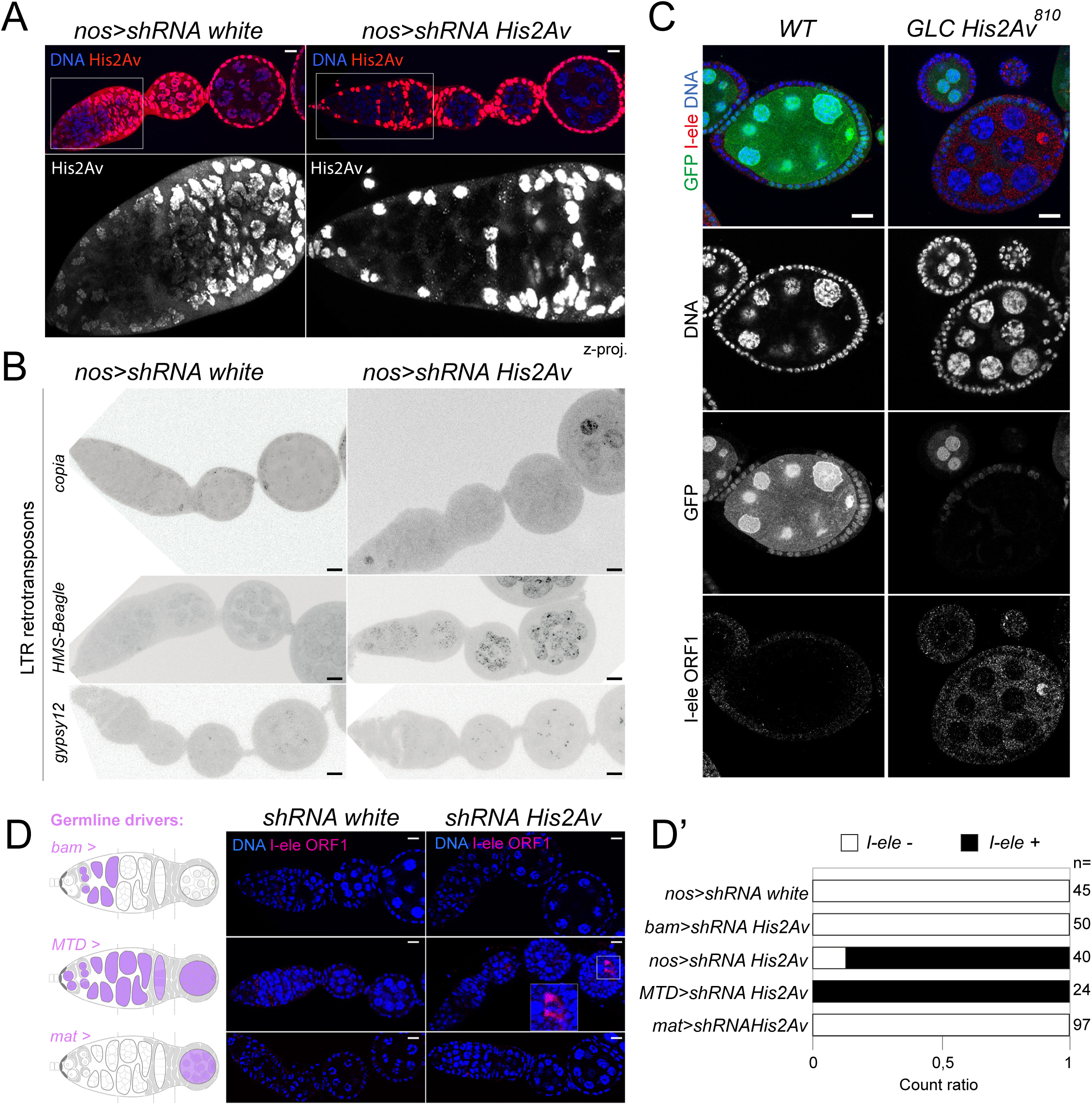
related to Fig.1. His2Av is required for TEs silencing. **A.** His2Av immunostaining in control *nos>shRNA white* and *His2Av* germline mutant ovaries. A magnification image shows the germarium. Scale bar: 10 µm. Z-proj: z projection. **B.** RNA FISH against transcripts from *copia*, *HMS-Beagle* and *gypsy-12* TEs in control *nos>shRNA white* ovaries and *His2Av* germline mutants. Scale bar: 10 µm. **C.** I-ele ORF1 immunostaining was done in Germline clones (GLC) mutant for *His2AV^810^* (absence of GFP). Scale bar: 10 µm. **D.** Scheme of *Drosophila* germarium and different germline drivers used in this study (violet). I-ele ORF1 immunostaining was done. A magnification image shows I-ele ORF1 accumulation in the oocyte in *MTD>shRNA His2Av* mutants. Scale bar: 10 µm. **D’.** I-element ORF1 quantification, present or absent in D.

**Supplementary Figure 2.**
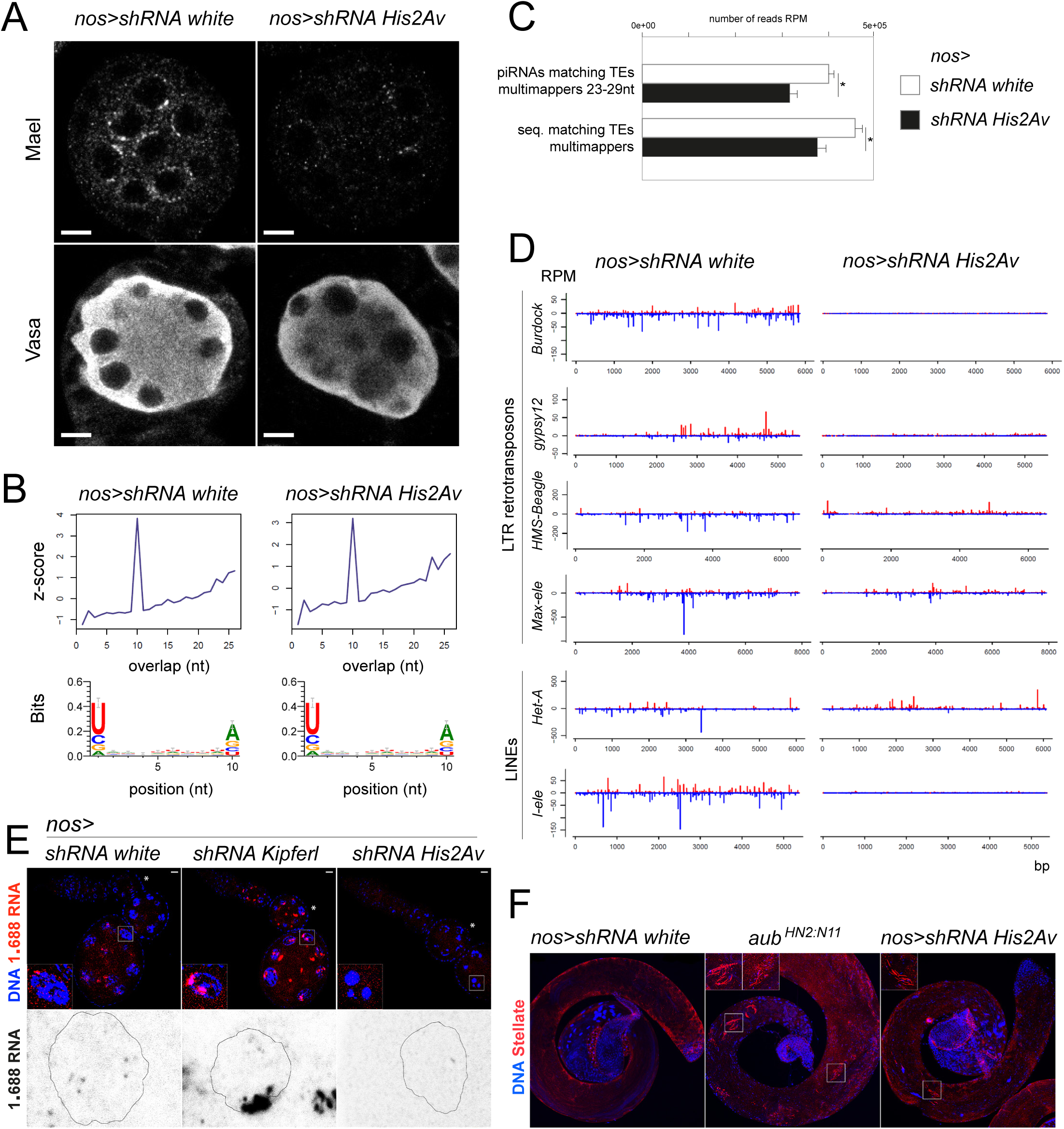
related to Fig.2. His2Av plays an important role in the piRNA pathway. **A.** Immunostaining of Mael and Vasa in *nos>shRNA white* and *His2Av* germline mutant ovaries. Scale bar: 10 µm. **B.** Ping pong signal calculated in multimappers piRNAs is shown in virgin *nos>shRNA white and His2Av* germline mutants. **C.** Multimapper piRNAs matching TEs on piRNA clusters defined in (Brennecke et al. 2007) in RPM are shown in virgin *nos>shRNA white and His2Av* germline mutants. **D.** Multimapping piRNAs (1 mismatch) originating from major TEs families in *nos>shRNA His2Av* mutants and control *nos>shRNA white*, in RPM. **E.** RNA FISH of DNA satellite 1.688 transcripts in *nos>shRNA white, kipferl and His2Av* germline mutants. Magnifications show Z-maximum intensity projection images of nuclei from stages 4-5. **F.** Testes from *white*, *aub^HN/N11^ and His2Av* germline mutants were fixed and stained for Stellate. Magnifications show the accumulation of Stellate crystals in mutant conditions.

**Supplementary Figure 3.**
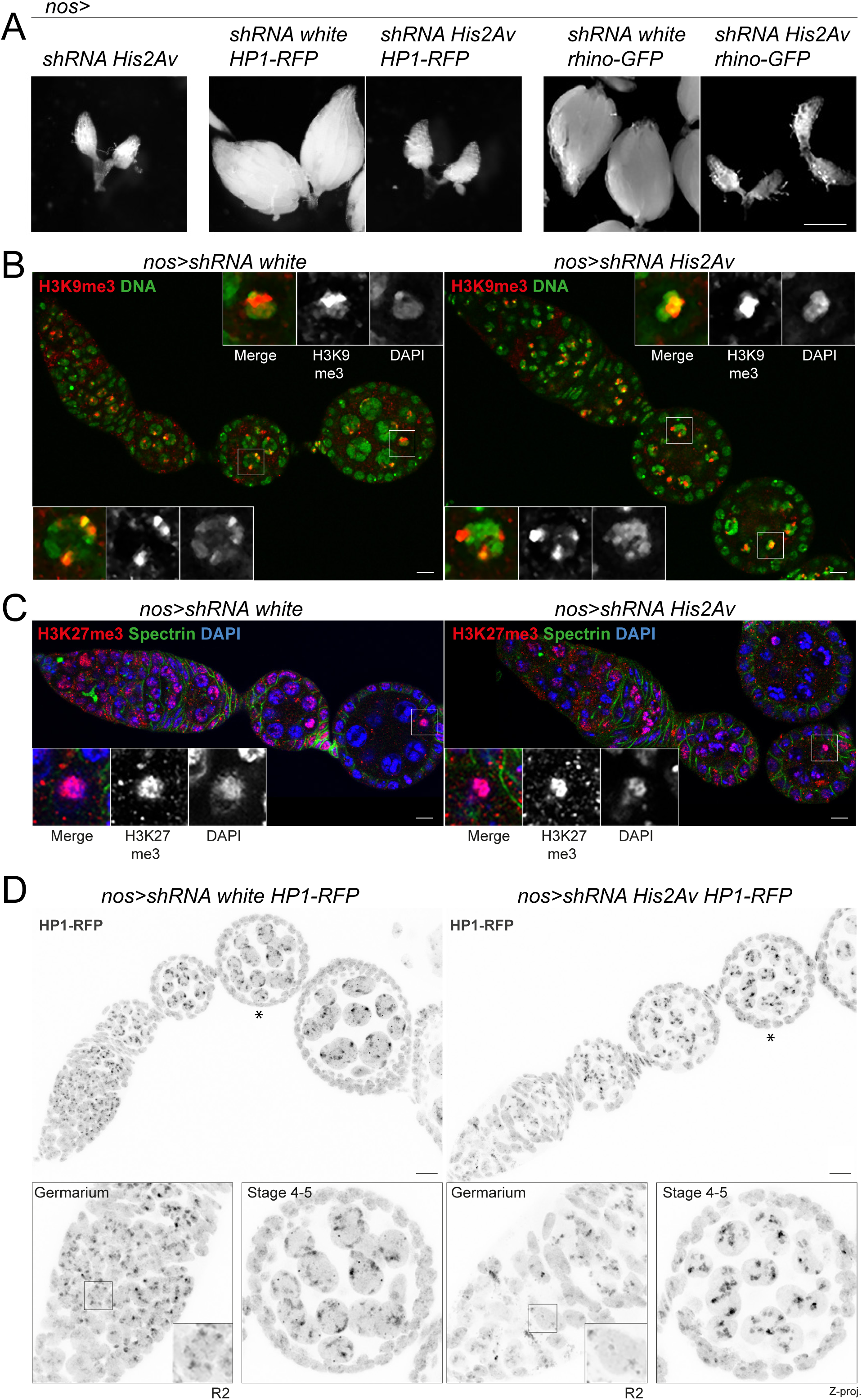
related to Fig.3. *His2Av* germline mutant flies accumulate oogenesis defects. **A.** Ovary size of *nos>shRNA white* or *His2Av* germline mutants expressing *HP1-RFP* or *rhino-GFP* are shown. **B.** H3K9me3 immunostaining in *nos>shRNA white* and *His2Av* germline mutant ovaries. Magnifications show the oocyte and a nurse cell. Scale bar: 10 µm. **C.** H3K27me3 immunostaining in *nos>shRNA white* and *His2Av* germline mutant ovaries. Magnifications show the oocyte. Scale bar: 10 µm. **D.** HP1-RFP signal is shown in control shRNA *white* and *His2Av* germline mutant ovaries. Magnifications show the germarium (R2, Region 2) and stages 4-5 (indicated by a star). Scale bar: 10 µm. Z-proj: z projection.

**Supplementary Figure 4.**
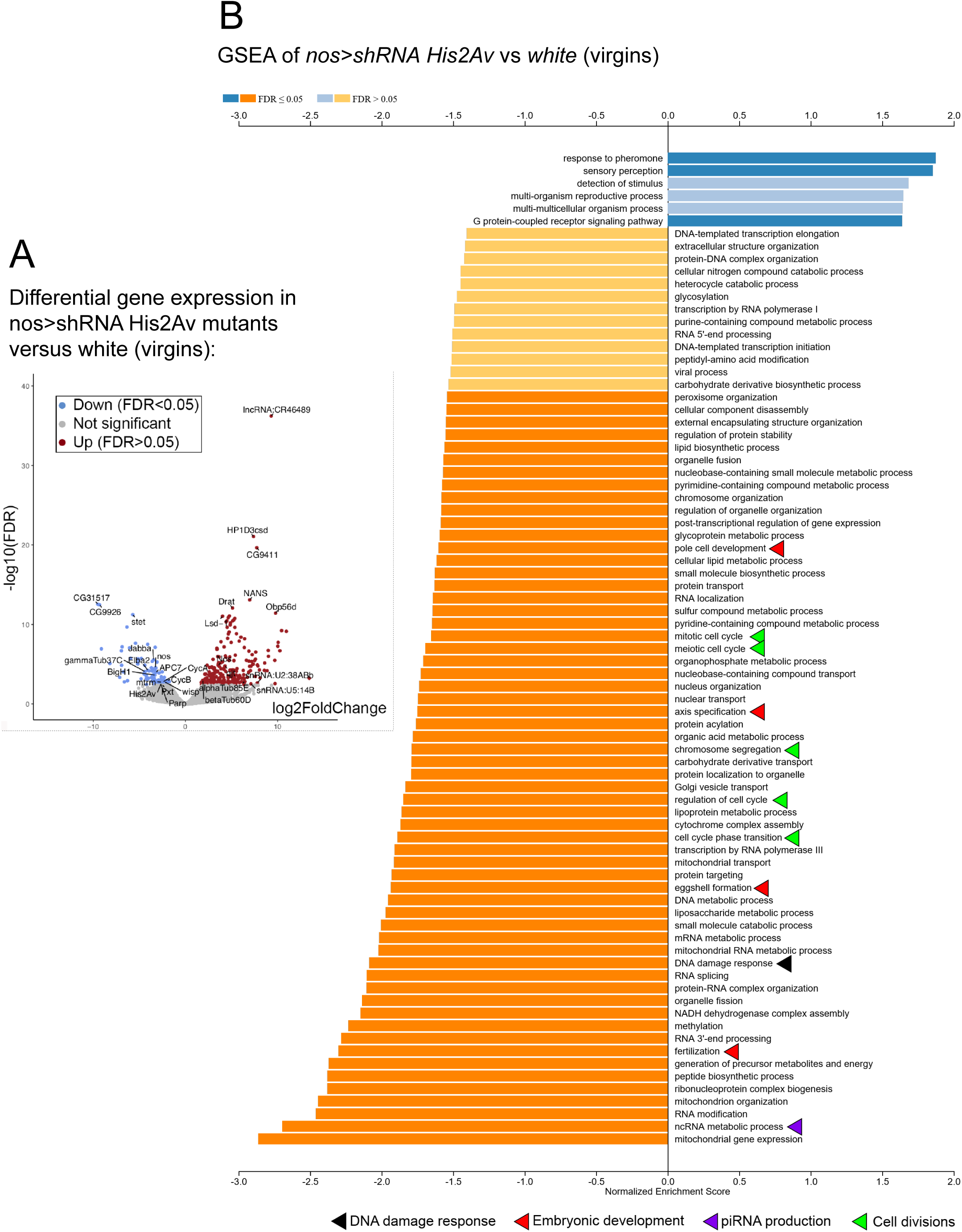
related to Fig.4. His2Av mutation leads to oogenesis arrest. **A.** Volcano plot showing genes differential expression in *nos>shRNA His2Av* germline mutant ovaries versus control *nos>shRNA white* ovaries, after DESeq2 analysis. Colored dots correspond to |log2FC|>2, FDR<0.05. Grey dots, not significant. **B.** Gene Set Enrichment Analysis (GSEA) of control *nos>shRNA white* versus *nos>shRNA His2Av* germline mutant ovaries transcriptomes performed on the WebGestalt webserver. Important oogenesis pathways are highlighted with colored arrows.

**Supplementary Figure 5.**
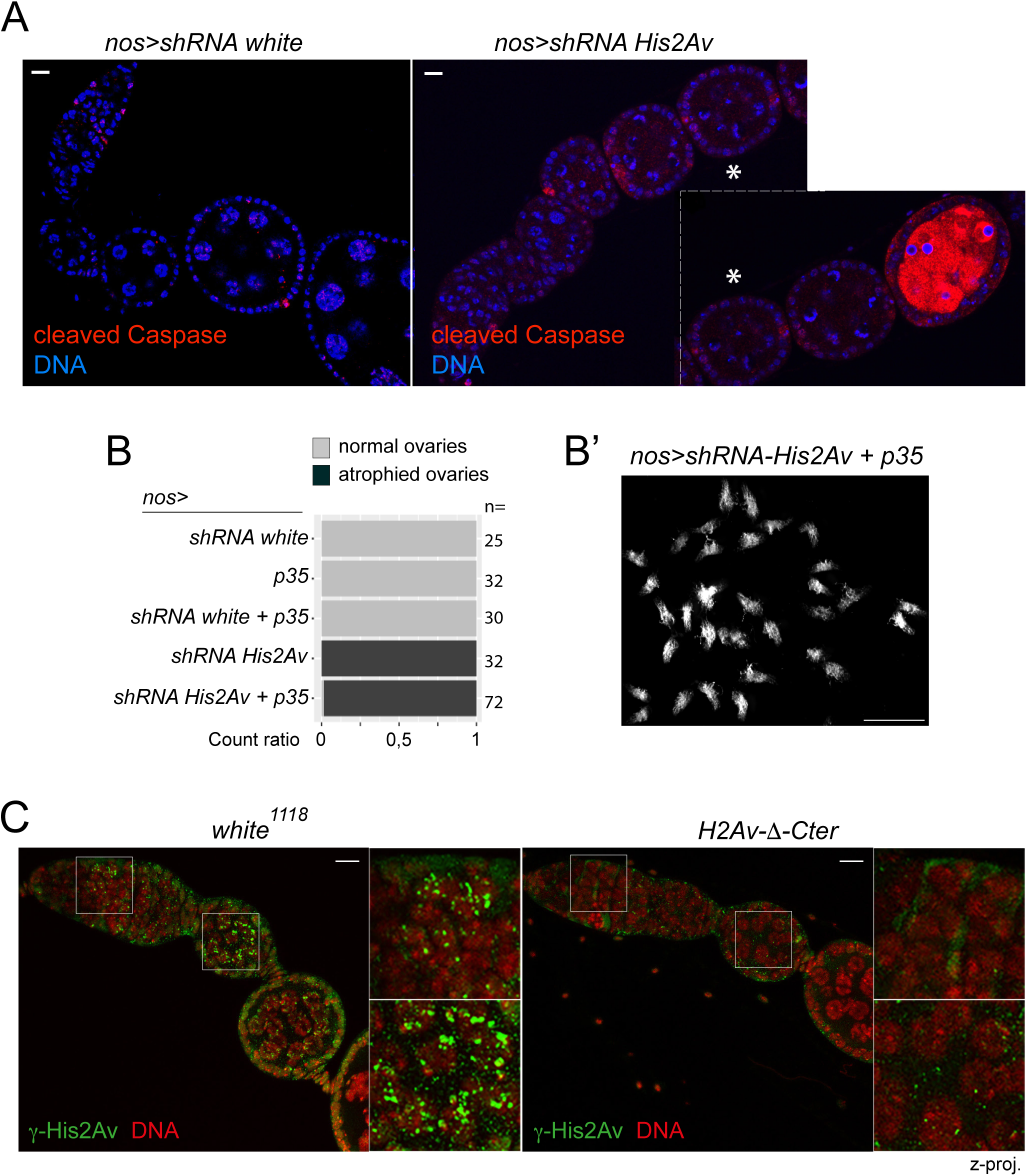
related to Fig.5. His2Av-Δ-Cter cannot be phosphorylated. **A.** Immunostaining of cleaved Caspase 3. Scale bar: 10 µm. The star shows the egg chamber that belongs to the same long arrested ovariole, separated by a dashed white line. **B.** Quantification of ovaries size upon over-expression of anti-apoptotic p35 at 22°C. **B’.** *nos>shRNA His2Av* germline mutant ovaries over-expressing anti-apoptotic p35 at 22°C. Scale bar: 2.5 mm. **C.** Phosphorylated γ-His2Av immunostaining in *nos>shRNA white* and *His2Av-Δ-Cter* germline ovaries. Magnifications show the meiotic region of the germarium and endoreplicating nurse cells. DNA is shown in red for better observation of γ-His2Av foci, in green. Scale bar: 10 µm. Z-proj: z projection.

**Supplementary Figure 6.**
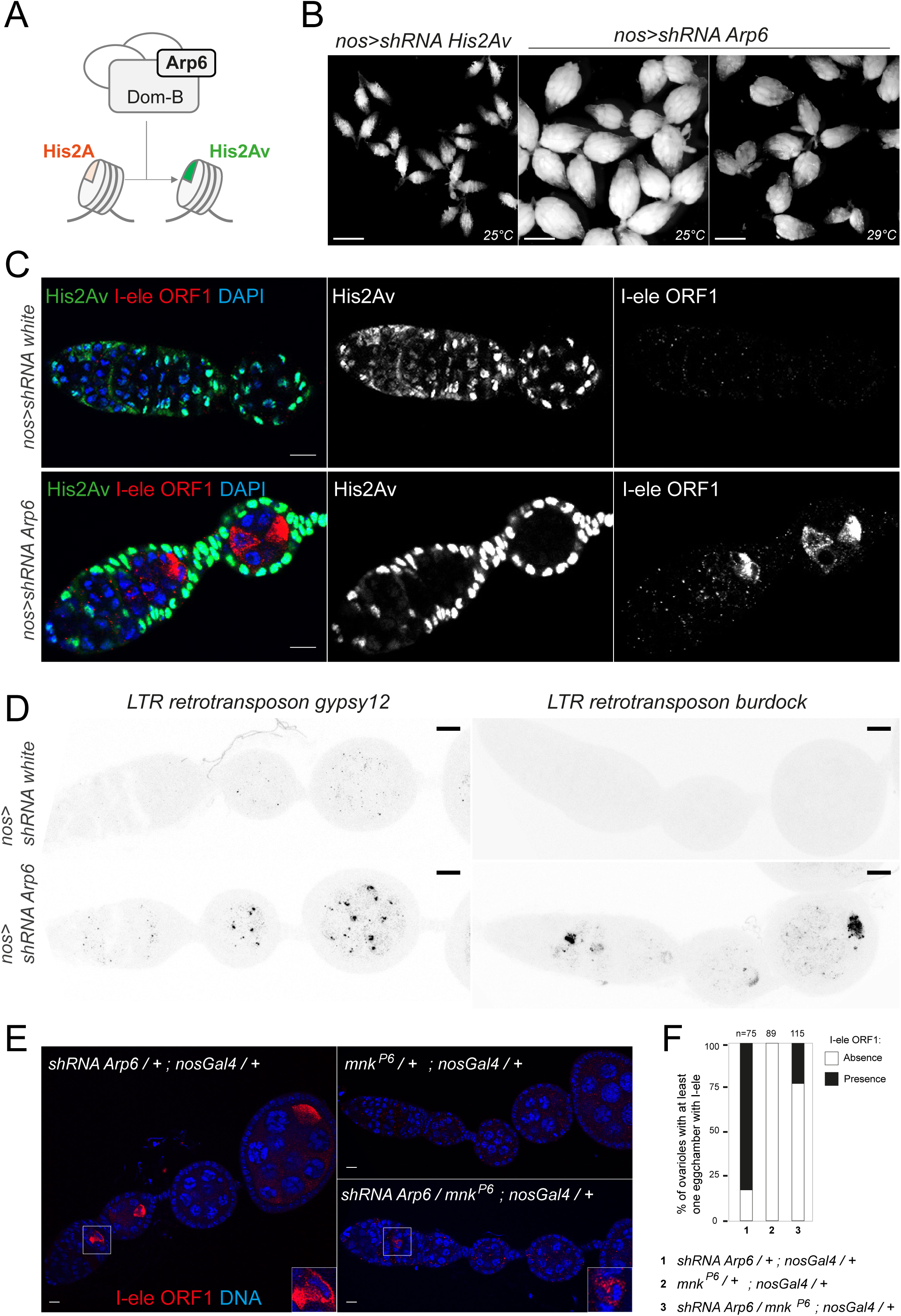
*Arp6* mutant ovaries recapitulate *His2Av* phenotype. **A.** Arp6/Dom-B complex responsible to include His2Av in the nucleosomes is depicted. **B.** Ovaries are shown in control *nos>shRNA His2Av* and *Arp6* germline mutants, at 25° or 29°C. **C.** Immunostaining in *nos>shRNA white* and *Arp6* germline mutants for I-ele and His2Av (25°C). Scale bar: 10 µm. **D.** RNA FISH against TEs transcripts from *gypsy-12* and *burdock* in *nos>shRNA white* and *Arp6* germline mutants. Scale bar: 10 µm. **E.** Immunostaining of I-ele ORF1 in *nos>shRNA Arp6* germline mutants, in heterozygous *chk2* (*mnk^P6^*) ovaries and in *nos>shRNA Arp6* and *chk2* (*mnk^P6^*) double mutants. **F.** Quantifications for E. The percentage of ovarioles with at least one egg chamber with I-ele ORF1 are shown for different genotypes.

**Supplementary Figure 7.**
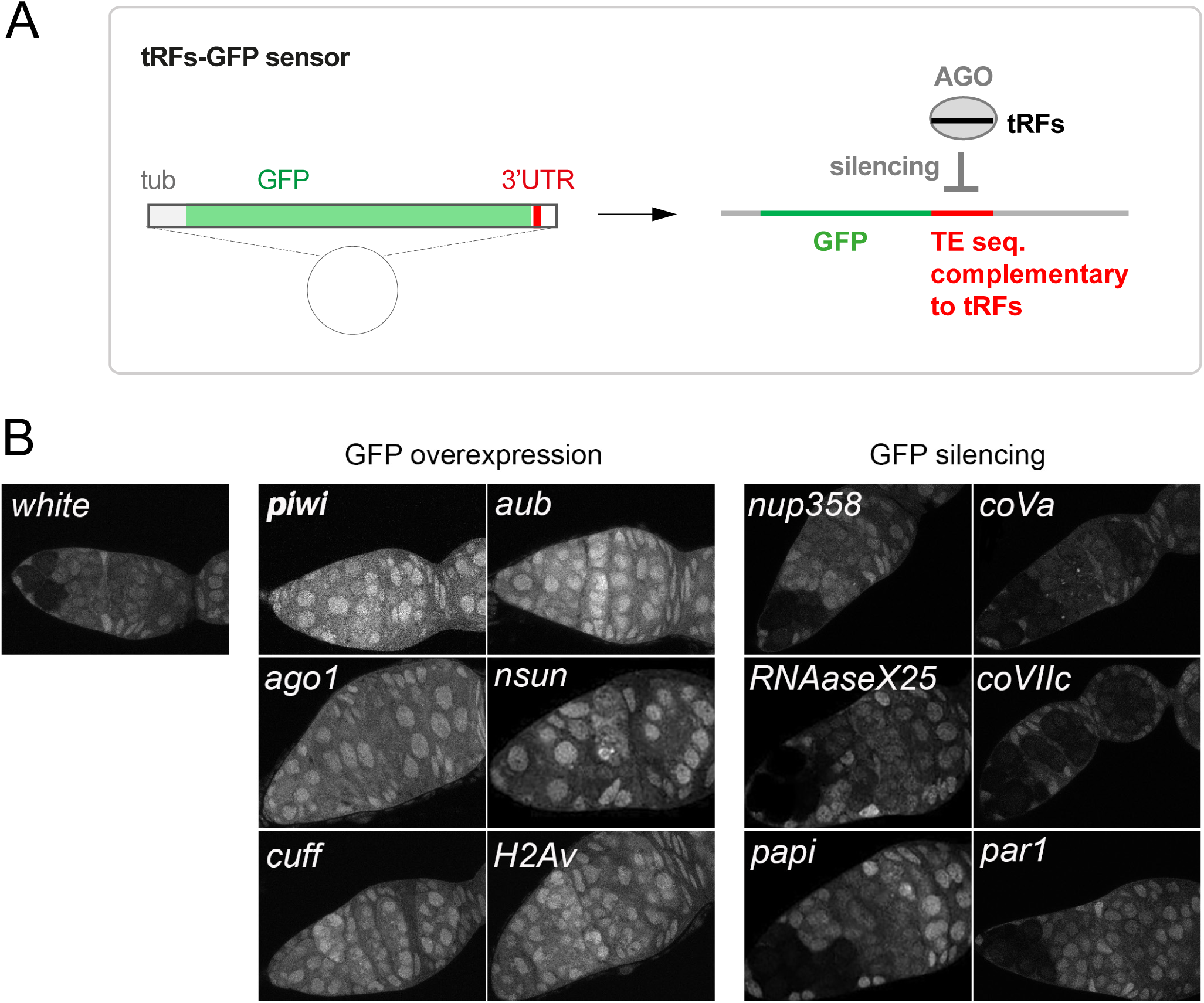
related to Material and Methods. tRFs-GFP sensor and genetic screen. **A.** Schematic of tRF-Lys-GFP sensor with a tubulin driver, followed of a GFP with a TE sequence complementary to tRFs-Lys inserted in the 3’UTR. **B.** tRFs-GFP signal in *nos>shRNA* control flies, mutants leading to a GFP overexpression in all germ cells, or mutants leading to an increased GFP silencing.

